# Coordinated chromosome motion emerges from mechanical coupling mediated by the physical spindle environment

**DOI:** 10.64898/2026.01.17.700110

**Authors:** Jiali Zhu, Kerry Bloom, Ehssan Nazockdast, Paul Maddox

## Abstract

During metaphase, chromosomes undergo oscillatory motion and exhibit distance-dependent coordinated movement with neighboring chromosomes within the spindle. However, the physical mechanism that gives rise to coordinated chromosome motion remains unclear. Here, we combine quantitative live-cell imaging in PTK1 cells, targeted perturbations of spindle microtubules and chromatin, and minimal mechanical modeling to uncover the mechanical basis of chromosome coordination during metaphase. We show that chromosome oscillations are dampened by stabilizing microtubules or by decondensing chromatin, yet inter-chromosomal coordination is preserved. The dissociation between chromosome oscillations and chromosome coordination suggests that coordination can arise from forces transmitted through the spindle environment. To test this idea, we developed a minimal mechanical model in which oscillating chromosome pairs are coupled through transient inter-chromosomal springs, while oscillations of each chromosome pair are generated by feedback between kinetochore-microtubule dynamics and centromere elasticity. The model demonstrates that stochastic mechanical connections are sufficient to generate correlated chromosome motion. To quantify chromosome coordination in a manner that reflects the mechanical properties of the surrounding spindle environment, we employed a microrheology-inspired analysis of time-lagged chromosome displacements. This framework reveals that microtubules and chromatin play distinct mechanical roles: microtubules primarily determine the spatial range and temporal build-up of coordination, while chromatin tunes its strength. Together, our results establish coordinated chromosome motion as an emergent mechanical property of the mitotic spindle, mediated by its viscoelastic properties and collective force transmission.

## Introduction

Accurate chromosome segregation relies on precise regulation of chromosome motion, which results from the interplay of dynamic spindle microtubules (Mitchison and Kirschner, 1984), mechanically compliant chromatin (Spicer and Gerlich, 2023), and kinetochores that serve as the physical interface between spindle microtubules and the chromosomes (Musacchio and Desai, 2017). Upon spindle interaction, chromosomes instantly start back-and-forth movements, which continue until the chromosomes undergo segregation (Skibbens et al., 1993; Burroughs et al., 2015). During metaphase, directional chromosome motion is coupled to the state of microtubule plus ends at kinetochores: depolymerization drives poleward movement, polymerization promotes anti-poleward motion (Asbury et al., 2006; DeLuca et al., 2006; Powers et al., 2009). Transitions between poleward and anti-poleward motion are regulated by tension across the centromere. As changes in centromere stretch modulate the mechanical load on sister kinetochores, microtubules switch between growth and shrinkage, thereby sustaining oscillatory chromosome motion during metaphase (Skibbens et al., 1995; Akiyoshi et al., 2010).

Chromosomes within the metaphase spindle also exhibit distance-dependent correlated motion, such that nearby chromosomes move more synchronously than those farther apart, and perturbations to spindle motors modulate the strength of this coordination (Vladimirou et al., 2013); however, other contributing factors remain unknown. One possible explanation is that the spindle itself coordinates the collective chromosome movements. This hypothesis is consistent with previous studies that show the mitotic spindle as an active viscoelastic gel, capable of transmitting forces and generating correlated motion among embedded objects (Brugúes and Needleman, 2014). The physical basis for such force transmission lies in the spindle’s structural composition: the microtubule network forms a dense, interconnected structure capable of transmitting and sustaining forces across micron length scales (Nazockdast and Redemann, 2020; Forth and Kapoor, 2017), while highly condensed chromatin exhibits intrinsic mechanical resistance and elastic properties, allow it to bear and respond to external forces (Man et al., 2021). Determining how forces are transmitted between chromosomes through this composite microtubule–chromatin environment is therefore essential for understanding the physical origin of chromosome coordination.

Here, we combine quantitative live-cell imaging, targeted perturbations, and minimal mechanical modeling to dissect the physical basis of coordinated chromosome motion during metaphase. We first establish baseline oscillatory behavior and inter-chromosomal coordination in control PTK1 cells. We then selectively perturb microtubule dynamics using low-dose nocodazole and reduce chromatin compaction via histone hyperacetylation, enabling us to examine how coordination responds to changes in spindle microtubule architecture and chromatin mechanics, respectively. To connect chromosome coordination to spindle mechanics, we analyze chromosome motion using a microrheology-inspired framework that treats each chromosome as an internal probe of the spindle’s viscoelastic properties. Correlations in chromosome displacements therefore report how mechanical interactions are transmitted and dissipated across the spindle. In parallel, we use a minimal mechanical model to test the extent to which coordinated chromosome motion can be explained by force balance alone, without invoking detailed molecular regulation. Together, our results show that chromosome coordination is an emergent mechanical property of the mitotic spindle, with microtubules and chromatin contributing in distinct yet complementary ways. This work reveals how the spindle’s viscoelastic architecture integrates local force generation with long-range mechanical coupling to maintain robust chromosome coordination during mitosis.

## Results

### Chromosomes show coordinated oscillatory movements during metaphase in PTK1 cells

We examined chromosome behavior in PTK1 cells stably expressing Hec1-GFP, a core outer kinetochore protein that binds to the plus ends of kinetochore microtubules. Using a standardized 3D time-lapse imaging workflow (Figure 1A), we recorded metaphase cells with clearly aligned chromosomes and well-defined spindle poles. Spindles were oriented horizontally so that kinetochore movements could be quantified along the x-axis. Fluorescent kinetochore puncta were tracked over time, and for each sister-kinetochore pair we computed its center-of-mass (CM) trajectory, which is used as a measurement of that chromosome pair’s motion (Figure 1B). Consistent with previous reports (Wan et al., 2012), CM trajectories revealed that chromosome pairs underwent continuous oscillations during metaphase (Figure 1D, top middle).

**Figure 1.**
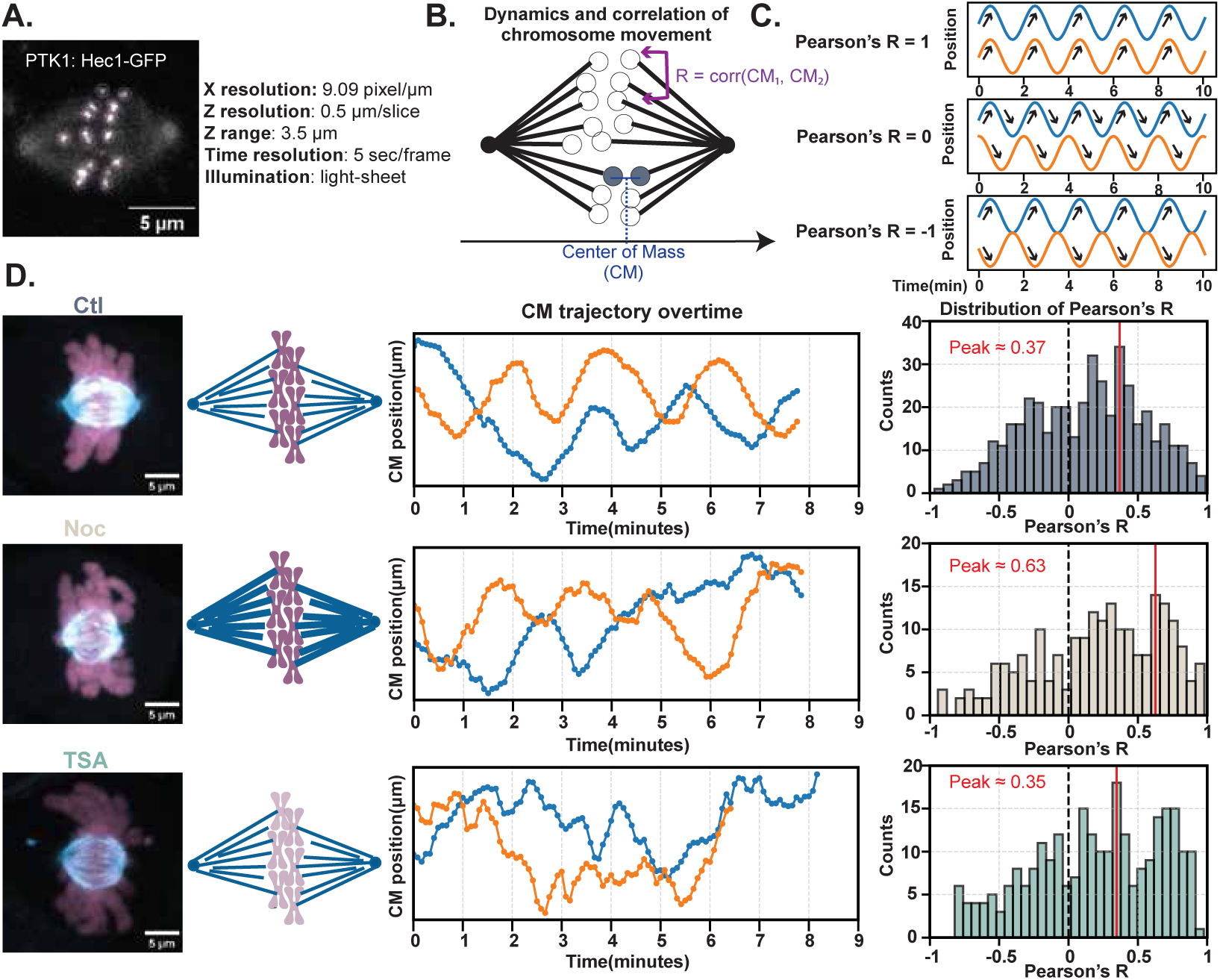
Metaphase chromosome oscillations are dampened after perturbing micro-tubule or chromatin structure, while coordination is maintained. **(A)** Representative metaphase PTK1 cells expressing Hec1-GFP, showing fluorescently labeled sister-kinetochore pairs used to track and quantify chromosome motions. Imaging parameters are listed. Scale bar, 5 *µm*. **(B)** Schematic illustrating the quantification of chromosome dynamics and coordination. For each spindle, the spindle poles were aligned horizontally, and motion was analyzed along the x-axis. The center of mass (CM) of each sister-kinetochore pair was defined as the mean x-position of the two kinetochores. Coordination between chromosomes was quantified by calculating Pearson’s correlation coefficients between CM trajectories. **(C)** Example trajectories illustrating different levels of correlation (Pearson’s R = 1, 0, −1). Arrows indicate the direction of displacement over time, highlighting synchronous, independent, or opposite-direction motion. **(D)** Left, maximum-intensity projections of spindle microtubules (blue) and chromosomes (magenta) in control (Ctl), nocodazole (Noc), and trichostatin A (TSA)–treated cells. Corresponding schematics depict relative changes in spindle microtubule organization and chromosome compaction. Middle, representative CM trajectories from two sister-kinetochore pairs over time. Control cells exhibit regular oscillatory motions, whereas both Noc– and TSA-treated cells show dampened chromosome movements. Right, distributions of Pearson’s correlation coefficients pooled from all CM pairs within individual spindles for control (N = 19 cells), nocodazole-treated (N = 13 cells), and TSA-treated (N = 15 cells) cells. The dashed line marks zero correlation, and the red line indicates the mode of each distribution. Despite reduced oscillation amplitude after Noc or TSA treatment, correlation distributions remain skewed toward positive values, indicating preserved chromosome coordination.

To determine whether different chromosome pairs move independently or in a coordinated manner, we quantified relationships between their CM trajectories by calculating Pearson’s correlation coefficients (Pearson’s R) for all CM pairs within each spindle (N = 19 cells) (Figure 1B). The CM for a chromosome pair is defined as the mean x-position of its two sister kinetochores. Pearson’s R ranges from –1 to 1 and reports how similarly two signals change over time (Figure 1C). Values near 1 reflect synchronous motion, values near –1 indicate motion in opposite directions, and values near 0 indicate no consistent relationship. In control cells, the distribution of Pearson’s R pooled across cells peaked at 0.35, indicating that different chromosome pairs exhibit positively correlated movements during metaphase (Figure 1D, top right).

These measurements established two baseline features of metaphase chromosome behavior in PTK1 cells: individual chromosome pairs undergo oscillatory motion over an 8-min time window as evidenced by the sinusoidal pattern of trajectories, and different chromosome pairs move in a coordinated manner as measured by Pearson’s R. Having defined these baseline properties, we next asked how they change when spindle or chromatin structure is perturbed.

### Chromosome dynamics are reduced but coordinated motion is maintained after perturbing spindle or chromatin structure

To determine how microtubules and chromatin influence coordinated chromosome motion, we perturbed these two structures pharmacologically in PTK1 cells. Nocodazole binds to *β*-tubulin, a subunit of the microtubule polymer, and prevents tubulin from assembling into microtubules. At high concentrations (1-10 *µ*M), nocodazole causes net depolymerization of the microtubule network (Laisne et al., 2021), whereas at low concentrations (4-100 nM), nocodazole primarily suppresses microtubule dynamics without substantial depolymerization (Jordan et al., 1992). Consistent with this regime, previous work in U2OS and HeLa cells has shown that 20 nM nocodazole stabilizes spindle microtubules, increases chromosome congression errors, and delays mitosis, while still allowing a substantial fraction of cells to proceed through mitosis (Rajendraprasad et al., 2021).

Guided by these established effects, we treated PTK1 cells with nocodazole (Noc) at 10 ng/mL (∼33 nM) for 1 h to selectively disrupt microtubule dynamics without abolishing spindle structure. Under these conditions, the spindle morphology remained intact, and many cells progressed into anaphase (Figure S1A). Quantitative analysis revealed that Noc increased spindle microtubule intensity by ∼23%, from 656 to 809 A.U./voxel (*p* = 0.038), while reducing spindle area by ∼27%, from 5,485 to 4,012 px^2^ (*p <* 0.001) (Figure S1C). In contrast, chromosome intensity and chromosome area were not significantly altered by Noc, remaining at 325 vs. 333 A.U./voxel (*p* = 0.92) and 12,715 vs. 13,460 px^2^ (*p* = 0.51), respectively (Figure S1D). These results indicated that low-dose Noc selectively perturbed spindle architecture without affecting chromatin compaction. Increased spindle intensity accompanied by reduced spindle area suggested a more compact and denser spindle.

Histone post-translational modifications (PTMs) actively regulate mitotic chromosome condensation. Deacetylation of histone H4 at lysine 16 enhances nucleosome–nucleosome interactions and promotes chromatin compaction during mitotic entry (Wilkins et al., 2014). To perturb mitotic chromatin compaction, we treated PTK1 cells with the pan-histone deacetylase inhibitor trichostatin A (TSA). Among the tested conditions, treatment with 500 nM TSA for 3 h produced the strongest increase in histone acetylation (Figure S1B) and was used for subsequent analyses. Hyperacetylation induced by TSA resulted in pronounced chromatin decondensation. Chromosome fluorescence intensity decreased by ∼26%, from 325 to 239 A.U./voxel (*p <* 0.001), accompanied by a ∼27% increase in chromosome area, from 12,715 to 16,085 px^2^ (*p <* 0.001) (Figure S1D). TSA did not significantly alter spindle organization: spindle area remained unchanged (5,485 vs. 5,614 px^2^, *p* = 0.85), and spindle microtubule intensity showed a modest, non-significant increase (from 656 to 744 A.U./voxel, *p* = 0.31) (Figure S1C).

Both Noc and TSA treatments altered chromosome motion from regular oscillations to more stochastic fluctuations over the 8-min time window (Figure 1D, middle). In control cells, chromosome center-of-mass (CM) trajectories exhibited continuous, near-sinusoidal oscillations, while both Noc– and TSA-treated cells showed reduced amplitudes, irregular fluctuations in chromosome CM positions. Despite this change in chromosome motion, Pearson’s R distributions remained positively skewed in all conditions. Notably, the distribution peak was highest in Noc-treated cells (0.63, *N* = 13), while TSA-treated cells (0.35, *N* = 15) closely matched control cells (0.36, *N* = 19) (Figure 1D, left).

Together, these results demonstrated that although chromosome motion became dampened after disruption of either spindle or chromatin organization, inter-chromosomal coordination was preserved. This dissociation between oscillatory behavior and coordination ruled out the possibility that coordinated chromosome motion results from synchronization of oscillations.

### Minimal mechanical model drives continuous chromosome oscillations through feedback between KT–MT dynamics and centromere tension

To identify the minimal mechanical components sufficient to generate and transmit chromosome motion, we developed a coarse-grained model of a single oscillating chromosome that serves as a baseline for extending the model to multiple chromosomes. In this model, chromosome motion is driven by force balance at the kinetochore, with oscillations generated by load-dependent feedback between chromosome velocity and the changing number of kinetochore–microtubule (KT–MT) attachments (Figure 2A). Each sister kinetochore is represented as a point mass connected by an elastic centromeric spring and coupled to dynamic microtubule ends through spring-like kinetochore-microtubule attachments. As the chromosome moves poleward, increased KT–MT attachment at the leading kinetochore amplifies poleward force, stretching the centromeric spring and generating a restoring force that eventually opposes and reverses motion, shifting pulling force to the opposite kinetochore. This feedback among force generation, KT-MT attachment number, and centromere elasticity produces self-sustained oscillations without invoking detailed molecular regulation.

**Figure 2.**
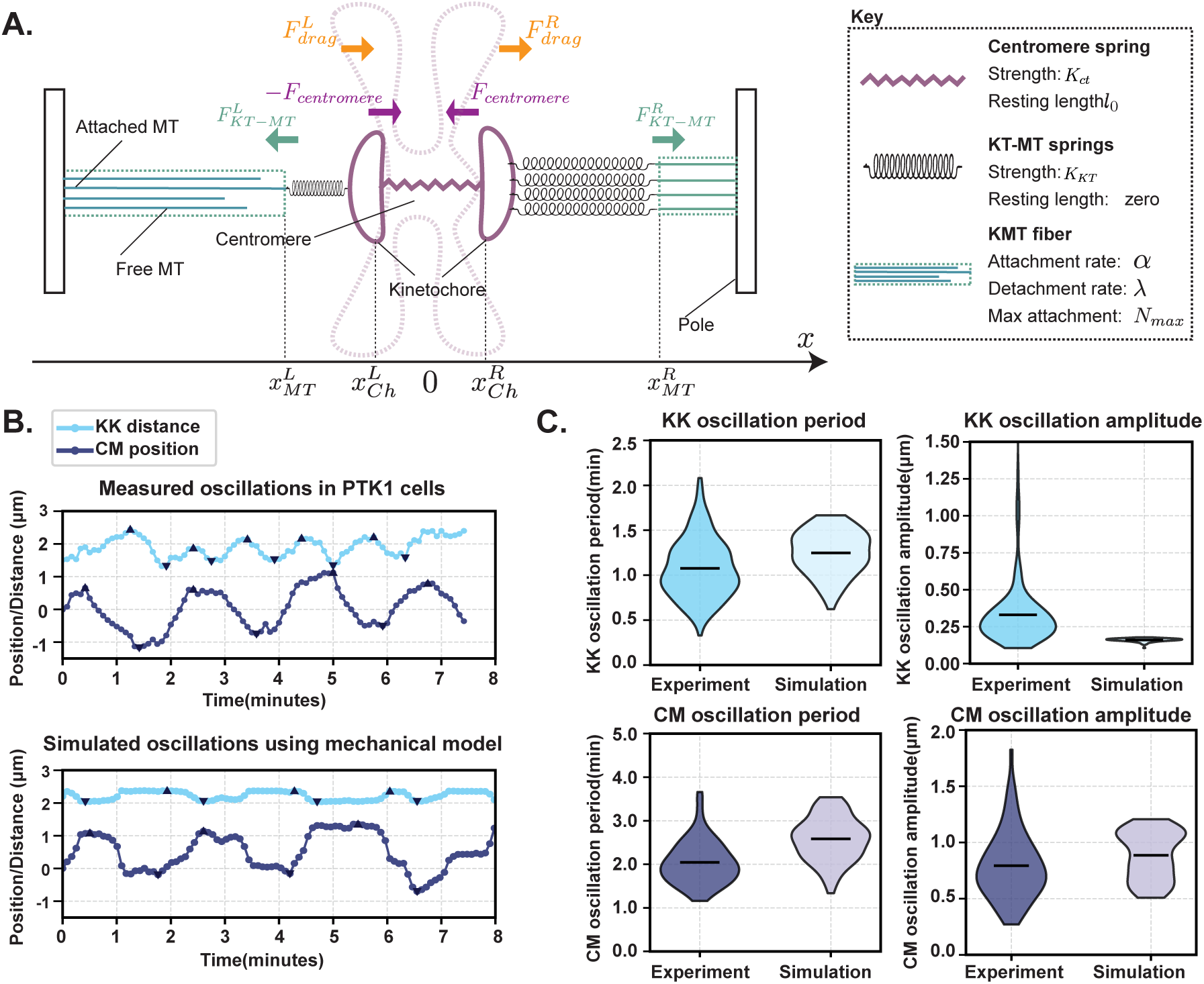
A minimal mechanical model recapitulates chromosome oscillations observed in PTK1 cells. **(A)** Schematic of the mechanical model used to simulate chromosome oscillations. Sister kinetochores are mechanically coupled by a centromeric spring (purple) with stiffness *K_ct_*and resting length *l*_0_. Each kinetochore interacts with dynamic microtubule plus ends via elastic kinetochore–microtubule (KT–MT) springs (black), whose number reflects the instantaneous number of attached microtubules. KT-MT attachments form and detach stochastically with rates *α* and *λ*, up to a maximum attachment number *N_max_*. Chromosome motion occurs along the spindle axis (x-axis) and is opposed by viscous drag within the mitotic spindle. Key model elements and parameters are summarized in the legend. **(B)** Representative oscillatory trajectories measured in PTK1 cells (top) and generated by the mechanical model (bottom). Two distinct oscillatory modes are shown: oscillation of the center of mass (CM) of the sister-kinetochore pair (purple) and oscillation of the inter-kinetochore distance (KK; blue). The model reproduces the characteristic frequency, phase relationship, and qualitative features of both CM and KK oscillations observed experimentally. **(C)** Quantitative comparison of oscillation period (left) and amplitude (right) for KK (top row, blue) and CM (bottom row, purple) oscillations measured experimentally and in simulations. Distributions are shown as violin plots; horizontal bars indicate median values. The model captures the characteristic periods of both CM and KK oscillations and reproduces the relative amplitude of CM oscillations, while underestimating the amplitude of KK oscillations compared with experiment.

The single-chromosome model generates oscillatory trajectories for both the chromosome-pair center of mass (CM) and the inter-sister kinetochore distance (KK), producing time series that resemble experimentally observed dynamics in PTK1 cells (Figure 2B). In both experiment and simulation, CM motion and KK distance oscillate with a stable phase relationship, with KK oscillations occurring at approximately twice the frequency of CM motion. The corresponding mean periods are 2.0 min for CM and 1.0 min for KK in experiment, and 2.6 min for CM and 1.25 min for KK in simulation.

To quantitatively compare model output with experimental measurements, we analyzed the distributions of oscillation periods and amplitudes for both KK and CM motion (Figure 2C). For KK oscillations, simulated periods overlapped with the experimental range but were shifted toward larger values, with mean periods of 1.25 min in simulation compared to 1.0 min in experiment. The model produced periodic centromere stretch with a lower mean amplitude and reduced variability relative to experiment, with mean amplitudes of 0.25 *µm* in simulation and 0.35 *µm* in experiment. For CM oscillations, simulated periods were similarly shifted toward larger values, with mean periods of 2.6 min in simulation compared to 2.0 min in experiment, while exhibiting a comparable overall spread. Simulated CM amplitudes fell within the experimental range, with mean amplitudes of 0.8 *µm* in simulation and 0.9 *µm* in experiment, although the simulated distribution was narrower. Variability in the simulated trajectories arose from stochastic diffusive forces on the chromosome applied at each time step and from stochastic fluctuations in kinetochore–microtubule attachment dynamics, which together generated the spread observed in the simulated period and amplitude distributions.

The model did not reproduce experimental measurements on a one-to-one basis, as expected given its deliberately simplified, coarse-grained design. Nevertheless, it captured the key qualitative and quantitative features of chromosome oscillatory dynamics relevant for analyzing inter-chromosomal mechanical relationships.

### A balance between centromere stiffness and KT–MT attachment dynamics ex-plains reduced chromosome activity

The single-chromosome model operates at the same scale as our experimental perturbations. We used the model to examine how mechanical parameters governing chromosome activity, focusing on two parameters: the centromere stiffness *K_ct_* (pN *µ*m^−1^) and the kinetochore–microtubule (KT–MT) attachment dynamics (*α* (s^−1^)). These parameters map directly onto our experimental treatments. Trichostatin A (TSA) treatment is expected to reduce effective centromere stiffness by decondensing chromatin, whereas nocodazole (Noc) treatment suppresses KT–MT attachment dynamics by dampening microtubule turnover.

To quantify chromosome activity, we computed the mean-squared velocity (MSV), defined as the time-averaged sum of the squared instantaneous velocities of the two sister kinetochores (Figure 3A). In simulations, MSV provides a scalar measure of overall chromosome activity: low MSV values correspond to trajectories dominated by small stochastic fluctuations in chromosome position, whereas high MSV values reflect highly dynamics chromosome oscillations (Figure 3A). In experiments, we measured MSV for individual chromosome pairs across multiple cells. In control cells, the average MSV was 5.61 ± 2.71 *µ*m^2^ min^−2^. MSV decreased to 4.59 ± 2.29 *µ*m^2^ min^−2^ following Noc treatment (P = 0.008) and further to 3.49 ± 1.76 *µ*m^2^ min^−2^ following TSA treatment (*P <* 0.001) (Figure 3B). Thus, oscillatory chromosome activity was reduced under both perturbations, with the most pronounced decrease observed following TSA treatment.

**Figure 3.**
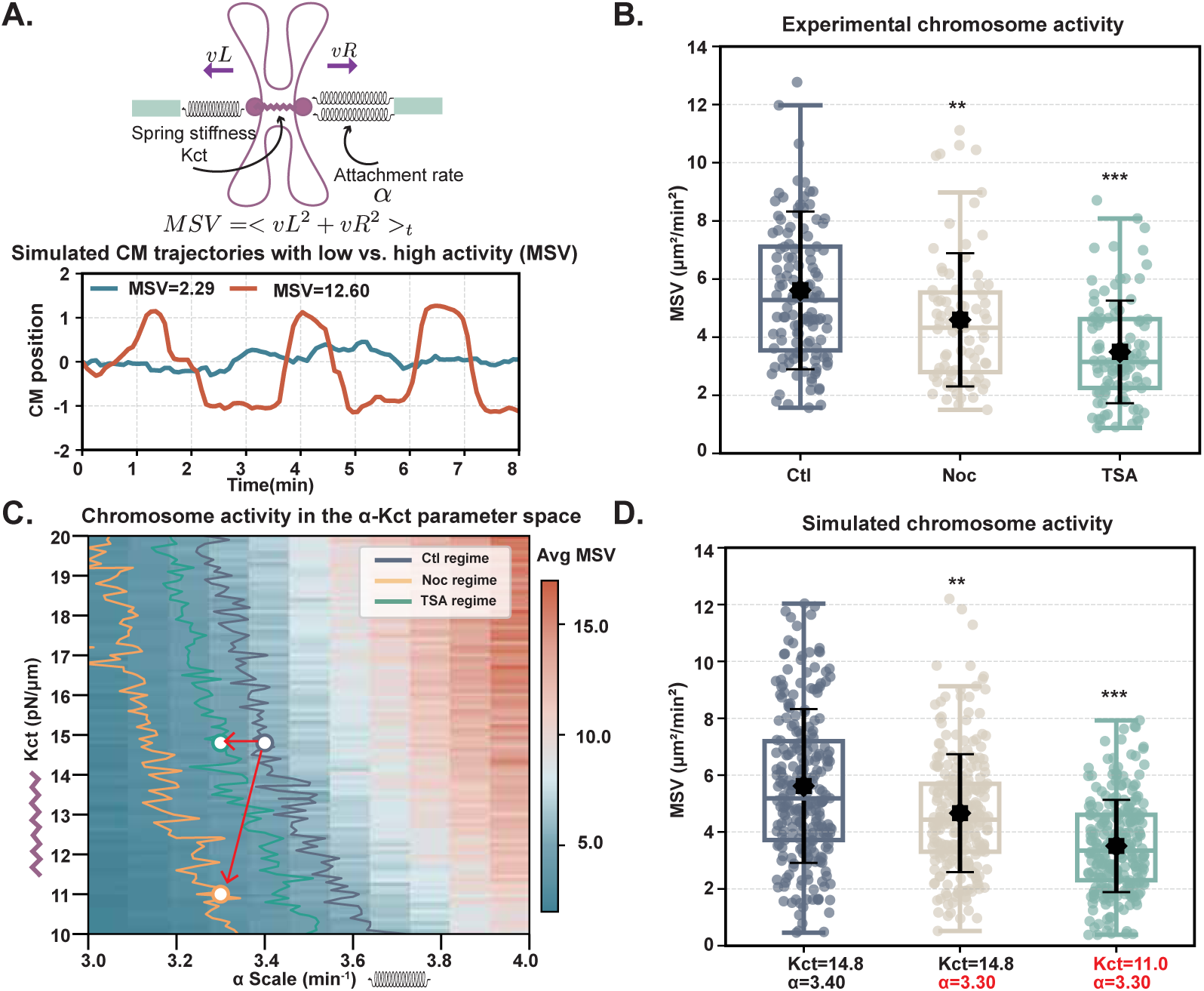
Chromosome activity reflects a balance between centromere stiffness and kinetochore-microtubule dynamics. **(A)** Schematic illustrating how chromosome activity was quantified using the mean squared velocity (MSV) of sister kinetochores. MSV integrates the instantaneous velocities of the left and right kinetochores over time, providing a single measure of overall chromosome motility. Bottom, representative simulated center-of-mass (CM) trajectories for chromosomes with low activity (blue, MSV = 2.29) and high activity (orange, MSV = 12.60), illustrating how MSV captures differences in the magnitude of chromosome movements. **(B)** Experimentally measured chromosome activity under control (Ctl), nocodazole (Noc), and trichostatin A (TSA) treatments. Each point represents one sister-kinetochore pair; boxplots summarize the distribution, and black symbols indicate the mean ± SD. Chromosome activity is significantly reduced after both Noc and TSA treatments. **(C)** Simulated chromosome activity across the two-parameter space of centromere stiffness (*K_ct_*) and kinetochore–microtubule attachment rate (*α*). The heat map shows the average MSV, and colored contours delineate regions corresponding to the experimentally measured MSV ranges for Ctl, Noc, and TSA conditions. White markers indicate representative parameter combinations selected for simulations shown in (D). Red arrows indicate shifts in *K_ct_* and *α* relative to the control regime. **(D)** Simulated MSV distributions for the parameter sets highlighted in (C). Each point represents one simulated chromosome pair; boxplots summarize the distributions, and black symbols indicate the mean ± SD. Simulation noise was tuned to match experimental variability. The model reproduces the relative decrease in chromosome activity observed experimentally following Noc and TSA treatments.

To distinguish how centromere stiffness (*K_ct_*) and KT–MT attachment dynamics (*α*) differentially shape chromosome activity (MSV), we computed MSV across the two-dimensional parameter space (*α, K_ct_*). We first analyzed the model in the absence of noise to establish the baseline deterministic behavior of the system. Without noise, the model produced a deterministic MSV landscape across *K_ct_*and *α* that revealed a sharp boundary separating non-oscillatory and oscillatory regimes (Figure S2D). At low *α* values (*<* 6*s*^−1^), where KT-MT attachments turned over slowly, oscillations occurred only when the centromere spring was relatively soft. As *α* value increased, faster KT-MT turnover required progressively stiffer centromere springs to sustain oscillations. It is likely that when the centromere is too stiff relative to slow KT–MT dynamics, restoring forces from the centromere springs dominate, dampening chromosome motion and preventing oscillations. Conversely, when KT–MT turnover is faster relative to a soft centromere, small changes in chromosome position can rapidly increase the number of attached microtubules, producing large, rapidly varying pulling forces that the centromere cannot buffer, leading to irregular, unstable motion. Thus, sustained oscillations occur only in a regime where centromere stiffness and KT–MT turnover are tuned to balance one another.

Introducing noise into the model (diffusive motion of chromosomes and KT–MT attachment fluctuations) transformed the deterministic chromosome activity phase diagram into a more continuous landscape (Figure S2D). Compared with the sharp boundary separating non-oscillatory and oscillatory regimes in the noise-free case, stochasticity smoothed the transition between regimes and shifted the parameter values at which changes in chromosome activity occur. In the noise-driven MSV heatmap (Figure 3C), higher centromere stiffness (*K_ct_*) shifted the onset of elevated chromosome activity to lower values of the KT–MT turnover rate (*α*). In other words, stiffer centromeres require slower microtubule dynamics and softer centromeres require faster dynamics to reach comparable activity levels. Elevated chromosome activity (MSV) therefore arises only within a narrow regime where centromere stiffness and KT–MT attachment dynamics are appropriately matched.

To link the model to experiment, we identified the regions of parameter space that reproduced the experimentally measured chromosome activity (MSV) values for each condition using contour lines (Figure 3C). For all three conditions, these contours spanned a narrow range of *α* values but extended across a much broader range of *K_ct_* values, indicating that chromosome activity (MSV) was far more sensitive to KT–MT dynamics than to centromere stiffness. By averaging each contour region, we extracted representative parameter values for control (*α* = 3.40 s^−1^*, K_ct_* = 14.8 pN *µ*m^−1^), Noc-treated (*α* = 3.30 s^−1^*, K_ct_* = 14.8 pN *µ*m^−1^), and TSA-treated (*α* = 3.30 s^−1^*, K_ct_* = 11 pN *µ*m^−1^) cells, which were shown as points along the contours (Figure 3C). For Noc treatment, the model indicated that a reduction in KT–MT dynamics (*α*) alone was sufficient to account for the observed decrease in chromosome activity, consistent with Noc’s known effect of stabilizing microtubules and suppressing KT–MT turnover. In contrast, TSA treatment shifted the system into a regime that required concurrent reductions in both KT–MT dynamics (*α*) and centromere stiffness (*K_ct_*). This behavior was consistent with chromatin decondensation leading to a softer, less effective centromere spring. Changes in KT–MT turnover may additionally have been associated with TSA’s effects on tubulin acetylation, as TSA is not a selective perturbation. In our experimental measurements, TSA treatment increased spindle intensity, although this effect did not reach statistical significance (Figure S1C). The model’s sensitivity to KT–MT turnover indicated that modest changes in microtubule dynamics were sufficient to alter chromosome activity, even when corresponding changes in spindle intensity were subtle.

### Dynamic inter-chromosomal springs provide a mechanism to generate and regulate coordination among chromosomes

We extended the single-chromosome model to include inter-chromosomal springs and examined the role of mechanical coupling between chromosomes in coordinating chromosome motion. In this extended model, the center of mass (CM) of each sister-kinetochore pair can stochastically connect to other CMs through springs characterized by a stiffness (*K_ic_*) and an effective length (*l_ic_*), which determines how far apart chromosomes can be while still forming transient mechanical connections (Figure 4A). These springs have zero mechanical rest length, such that no preferred inter-chromosomal spacing is imposed. Inter-chromosomal springs form and dissociate between CM pairs at rates *k_on_* and *k_off_*, respectively. When two CMs are connected, their relative displacement generates an additional coupling force that acts on top of the centromere restoring forces and KT–MT pulling forces. This stochastic network of inter-chromosomal connections enables force transmission between chromosomes, thereby generating correlated chromosome oscillations across the spindle.

**Figure 4.**
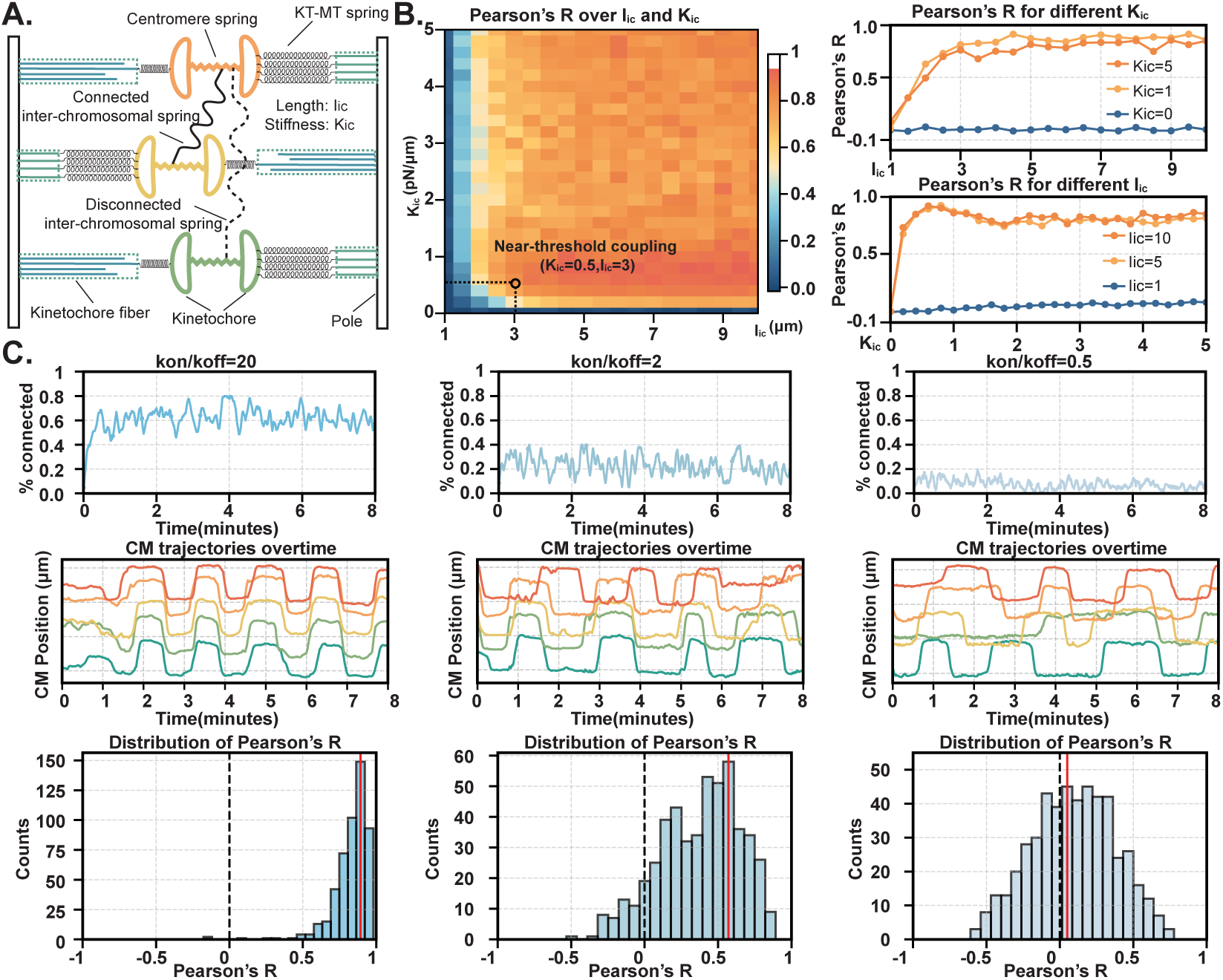
Dynamic inter-chromosomal springs generate coordinated chromosome motion, with correlation strength set by coupling dynamics. **(A)** Schematic illustrating the extension of the single-chromosome mechanical model (Figure 2) to a multi-chromosome system. The center of mass (CM) of each sister-kinetochore pair is coupled to neighboring chromosomes through stochastic inter-chromosomal springs. Each spring is characterized by a stiffness *K_ic_* and an effective length *l_ic_*, and dynamically forms and dissociates according to attachment and detachment rates. **(B)** Simulated Pearson’s correlation coefficient (R) as a function of inter-chromosomal spring stiffness and effective length. Left, heatmap showing the average correlation across the stiffness-length parameter space. Right, line plots illustrating that correlation saturates over a broad range of stiffness and effective length values and is relatively insensitive to further increases once mechanical coupling is established. We fixed inter-chromosomal spring stiffness and effective length at the minimum values that produced detectable, nonzero correlation (near-threshold coupling) when examining effects of other parameters. **(C)** Effects of inter-chromosomal spring dynamics on chromosome coordination. All simulations use identical spring mechanical properties (near-threshold coupling), while varying the ratio of attachment to detachment rates. Top row, fraction of chromosome pairs connected by inter-chromosomal springs over time,illustrating decreased connectivity with slower spring dynamics. Middle row, corresponding CM trajectories for five chromosomes, showing progressive loss of synchrony as spring dynamics decrease. Bottom row, distributions of Pearson’s correlation coefficients pooled across multiple simulation runs. Fast spring dynamics yield narrowly distributed, strongly positive correlations, whereas slower dynamics produce broad distributions centered near zero.

We examined how centromere stiffness (*K_ct_*) and KT–MT attachment dynamics (*α*), previously shown to regulate individual chromosome activity, shaped inter-chromosomal coordination in the multi-chromosome model. Using the same two-dimensional parameter sweep of centromere stiffness (*K_ct_* = 10–20 pN *µ*m^−1^) and KT–MT turnover rate (*α* = 3–15 s^−1^), we computed both chromosome activity, quantified by MSV, and coordination, quantified by Pearson’s R, in the presence of inter-chromosomal springs. Introducing mechanical coupling altered individual chromosome activity: across control-, Noc-, and TSA-mapped regimes, the combinations of KT–MT turnover and centromere stiffness that reproduced the experimentally measured MSV values were shifted toward higher KT–MT turnover rates relative to the uncoupled model (Figure S3A). Thus, in the presence of the inter-chromosomal springs, faster KT–MT attachment dynamics were required to sustain a given level of chromosome activity. At KT–MT turnover rates that reproduced the experimentally measured chromosome activity (MSV) in single chromosomal model, the presence of inter-chromosomal springs amplified stochastic diffusive fluctuations in individual chromosome motion, thereby suppressing coherent oscillatory behavior (Figure S3C).

In contrast to chromosome activity, chromosome coordination exhibited weaker sensitivity to changes in KT–MT attachment dynamics (*α*) and centromere stiffness (*K_ct_*). When the contours corresponding to the experimentally measured chromosome activity (MSV) in the presence of inter-chromosomal springs were overlaid onto the chromosome coordination heatmap, Pearson’s R remained relatively uniform along these contours (Figure S3B). This behavior mirrored the experimental observation that although Noc and TSA substantially suppressed chromosome activity, overall coordination between chromosomes was largely preserved. Because coordination was comparatively insensitive to *α* and *K_ct_*, we fixed these parameters at *α* = 6.2 s^−1^ and *K_ct_* = 12.3 pN *µ*m^−1^, which lay at the minimum of the deterministic chromosome activity (MSV) phase diagram where KT–MT dynamics and centromere stiffness were optimally balanced (Figure S2B). This choice allowed us to isolate the effects of inter-chromosomal coupling on coordination while holding individual chromosome activity constant.

To assess how inter-chromosomal mechanical coupling influenced coordination, we performed a two-parameter sweep over inter-chromosomal spring stiffness (*K_ic_*) and effective length (*l_ic_*). Across this parameter space, the average Pearson’s R between chromosome center-of-mass (CM) trajectories was largely insensitive to both *K_ic_* and *l_ic_* once coupling was established (Figure 4B). When the spring length was very short, Pearson’s R remained near zero because chromosomes could not effectively connect. As *l_ic_*increased to ∼3 *µ*m, Pearson’s R rose sharply but then remained approximately constant out to *l_ic_* = 10 *µ*m. Similarly, when spring stiffness was zero, Pearson’s R was near zero, whereas once stiffness exceeded a modest threshold (*K_ic_* = 1 pN *µ*m^−1^), correlation increased but showed little further dependence on stiffness between 1 and 5 pN *µ*m^−1^. Together, these results indicated that Pearson’s R robustly reported the presence of mechanical coupling but was not sensitive to the strength of coupling once a minimal threshold was crossed.

In contrast to inter-chromosomal spring stiffness (*K_ic_*) and effective length (*l_ic_*), the dynamics of inter-chromosomal coupling controlled the level of coordination by regulating the fraction of chromosomes that remained mechanically connected. At high connection stability (*k_on_/k_off_*= 20), most chromosomes were persistently linked, resulting in highly synchronized CM trajectories and a strongly positively skewed distribution of Pearson’s R (Figure 4C, left). At intermediate turnover rates (*k_on_/k_off_* = 2), transient coupling produced partially correlated oscillations and a broader distribution of Pearson’s R (Figure 4C, middle). When connections were infrequent (*k_on_/k_off_* = 0.5), chromosomes moved largely independently and the correlation distribution was centered near zero (Figure 4C, right).

Together, these results demonstrated that dynamic inter-chromosomal springs were sufficient to generate correlated chromosome motion, with the degree of coordination primarily controlled by the kinetics of spring formation and breakage. The physical properties of inter-chromosomal springs did not substantially alter the strength of coordination as measured by Pearson’s R.

### A two-point microrheology-inspired framework to study chromosome coordination across space and time

Pearson’s correlation summarizes coordinated motion by averaging over time and across chromosome pairs, which limits sensitivity to lag-dependent coordination and is limited by the number of chromosome pairs observed per cell. To address these limitations, we adopted a discrete, two-point microrheology–inspired framework based on time-lagged pairwise displacements, which increased the effective sample size and enabled coordination to be quantified within defined temporal windows (Figure 5A). Specifically, we tracked the center-of-mass (CM) trajectories of all sister-kinetochore pairs and computed their time-lagged displacements over a range of lag times *τ*, defined as Δ*X_i_*(*t, τ*) = CM*_i_*(*t* + *τ*) − CM*_i_*(*t*), where *t* is the absolute time and *τ* is the lag time. The smallest lag (*τ* = 5 s) corresponds to the imaging frame interval. For each unique CM pair (*i, j*) with *i* ≠ *j*, we quantified their coordination by calculating the ensemble-averaged product of their displacements:

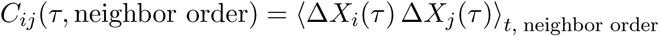

**Figure 5.**
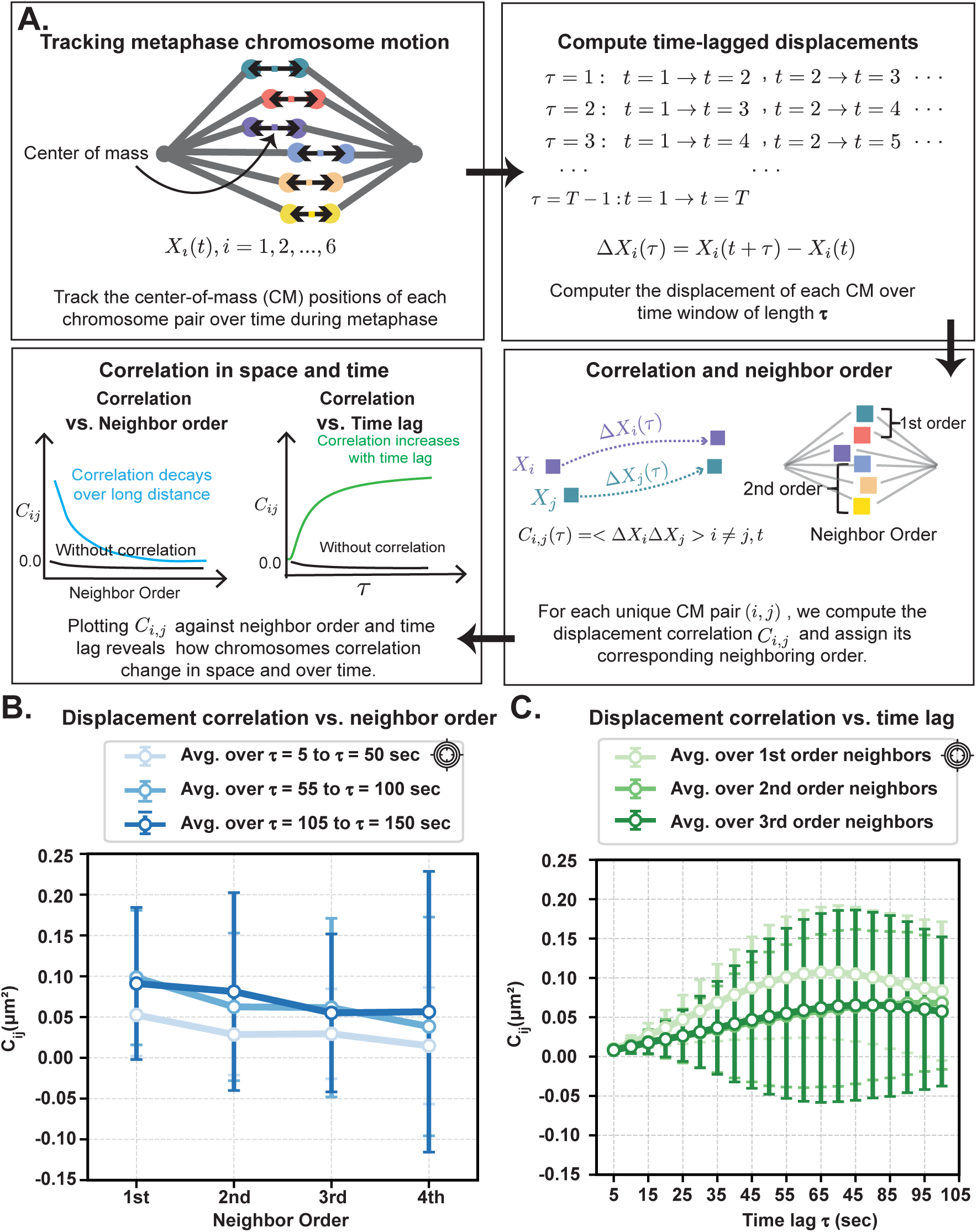
A two-point microrheology–inspired framework reveals that chromosome coordination reflects the viscoelastic properties of the mitotic spindle environment. **Figure 5. A two-point microrheology–inspired framework reveals that chromosome coordination reflects the viscoelastic properties of the mitotic spindle environment**. **(A)** Schematic overview of the time-lagged displacement correlation analysis used to quantify chromosome coordination across space and time. Step 1: The center-of-mass (CM) position of each sister-kinetochore pair is tracked throughout metaphase, yielding a set of CM trajectories *X_i_*(*t*), where *i* denotes the chromosome pair index ordered by spatial position within the spindle. Step 2: Time-lagged displacements Δ*X_i_*(*τ*) are computed for each trajectory, representing CM displacements over time windows of length *τ*. Time lags range from the temporal resolution of the data (5 s) up to the full trajectory duration. Step 3: Chromosome pairs are grouped by neighbor order based on their spatial separation along the spindle transverse (y) axis, with first-order neighbors defined as immediately adjacent CM pairs. For each chromosome pair (*i, j*) and time lag *τ*, a displacement correlation is computed and these correlations are ensemble-averaged over time and over all chromosome pairs belonging to the same neighbor order, yielding displacement correlation as functions of neighbor order and time lag. Step 4: The resulting displacement correlations depend on both spatial separation and time scale. For a viscoelastic spindle environment, correlations are expected to decrease with increasing neighbor order and to increase with longer time lags. **(B)** Displacement correlation as a function of neighbor order in control cells. CM trajectories were sampled at 5 s resolution. Correlations are averaged over three time-lag windows (5–50 s, 55–100 s, and 105–150 s). Correlation decreases with increasing neighbor order across all lag windows, with similar spatial dependence for the two longer-lag groups. Subsequent analyses therefore focus on spatial trends within the 5–50 s time-lag window. Lines and error bars indicate the mean ± SD. **(C)** Displacement correlation as a function of time lag in control cells. Correlations are averaged separately over first-, second-, and third-order neighbors. Correlation increases with time lag for all neighbor orders and approaches a plateau at longer time scales. Because the temporal dependence for second– and third-order neighbors is similar, subsequent analyses focus on the time-dependent correlation of first-order neighbors. Lines and error bars indicate the mean ± SD.

Here, the ensemble average was taken over time *t* and over all CM pairs belonging to the same neighbor order. Neighbor order served as a discrete proxy for inter-chromosomal distance, allowing us to characterize how coordination depended on spatial separation while avoiding confounding fluctuations in instantaneous distance between chromosomes.

This analysis is motivated by concepts from classical two-point microrheology, in which correlated motion of embedded tracer particles is used to infer the viscoelastic properties of the surrounding medium. The mitotic spindle exhibits both elastic, spring-like and viscous, fluid-like responses (Brugúes and Needleman, 2014), suggesting that chromosome motion may display analogous spatially and temporally structured correlations. In a viscoelastic medium, two general behaviors are expected. First, displacement correlations should decay with distance: perturbing one embedded particle strongly influences nearby particles but has diminishing influence on more distant ones. Second, displacement correlations should grow with time lag: at short timescales, motion is dominated by local fluctuations and correlations are weak, whereas at longer timescales, correlations associated with collective flows of the medium emerge (Figure 5A, last panel).

Using this framework, we examined how displacement correlations varied with both neighbor order (spatial dependence) and time lag *τ* (temporal evolution). We first applied this analysis to CM trajectories collected from 19 untreated PTK1 cells. Consistent with expectations for a viscoelastic medium, displacement correlation decreased with increasing neighbor order (Figure 5B). For clarity, data were grouped into three time-lag windows: 5–50 s, 55–100 s, and 105–150 s. Correlation curves from the two latter time windows largely overlapped, whereas the first time-lag group exhibited distinct behavior; Subsequent analyses were focused on the first time-lag window. We also examined how *C_ij_* evolved over time for first-, second-, and third-order neighbors (Figure 5C). Across all three neighbor orders, displacement correlation increased with time, consistent with the emergence of collective motion over longer timescales. Because second– and third-order neighbors exhibited largely overlapping trends, subsequent quantitative comparisons focused on first-order neighbors, which showed the strongest coordination.

### Inter-chromosomal spring length sets coordination range, while stiffness sets coordination strength

Having established a framework to quantify chromosome coordination across space and time, we used it to determine how coordination was altered by perturbations to microtubules and chromatin. We therefore compared how displacement correlations varied with both spatial separation and temporal lag following nocodazole (Noc) and trichostatin A (TSA) treatments. Given that coordination strength depends on the dynamics of inter-chromosomal spring connectivity, we fixed these dynamics at levels that matched experimentally observed coordination in control cells. This calibration allowed us to focus specifically on how inter-chromosomal spring properties shaped coordination. Specifically, using the displacement correlation of first-order neighbors at the 5-50 s lag window (first data point in the light blue curve, *C_ij_* ∼ 0.05; Figure 5B), we identified a region in the *k_on_*-*k_off_* parameter space where the simulated average correlation matched experimental levels (Figure S4A). Because multiple *k_on_*, *k_off_* combinations along this contour reproduced the same mean experiment-level correlation, we selected a representative point (*k_on_, koff*) = (20, 40) for all subsequent simulations. At these values, inter-chromosomal springs formed and broke on timescales that produced transient connectivity among ∼20-40% of chromosome pairs at any given time (Figure S4C).

Focusing on coordination across neighbor order within the 5–50 s time window (Figure 6A), we showed that in control cells, displacement correlation decayed with increasing neighbor order. This spatial decay became markedly shallower following both Noc and TSA treatment, indicating that correlated motion persisted over larger inter-chromosomal separations compared with control cells. Furthermore, the overall magnitude of coordination was selectively reduced in TSA-treated cells relative to both control and Noc-treated cells, reflecting weakened global coupling on top of its broader spatial reach after TSA treatment (Figure 6A). Thus, these perturbations induced two distinct effects on coordination:(i) a change in the spatial range of coordination (how far correlations extend across the spindle), and (ii) a change in the strength of coordination (the absolute magnitude of correlated motion).

**Figure 6.**
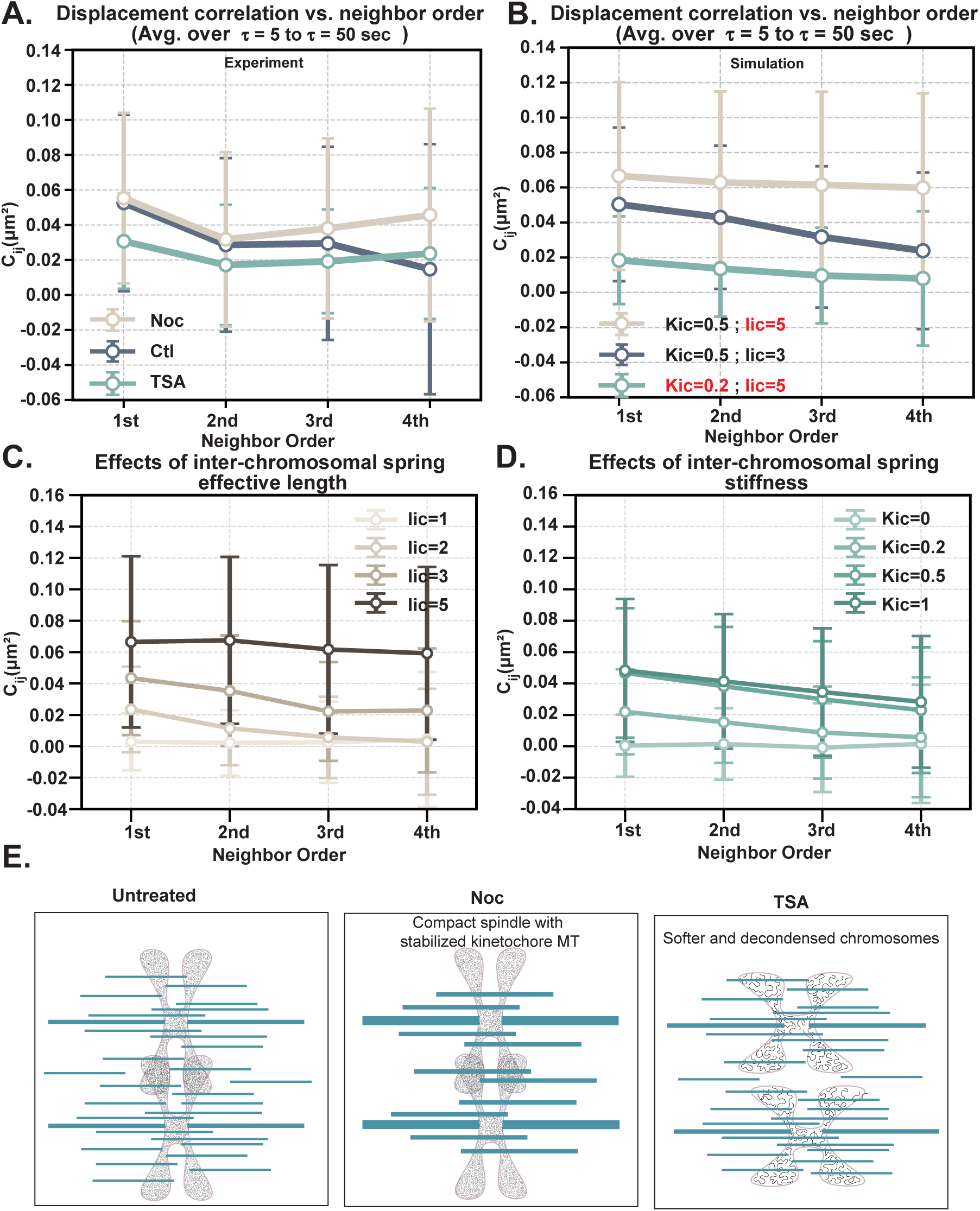
Inter-chromosomal spring length sets the spatial range of coordination, whereas spring stiffness controls coordination magnitude. **Figure 6. Inter-chromosomal spring length sets the spatial range of coordination, whereas spring stiffness controls coordination magnitude**. **(A)** Displacement correlation as a function of neighbor order for control (Ctl), nocodazole-treated (Noc), and trichostatin A–treated (TSA) cells. Correlations are averaged over time lags from 5–50 s. Relative to control, both Noc-and TSA-treated cells show higher correlations at larger neighbor orders. Across all neighbor orders, TSA-treated cells display lower mean correlation values than both control and Noc. Points and error bars indicate the mean ± SD. **(B)** Simulated displacement correlations as a function of neighbor order for selected combinations of inter-chromosomal spring stiffness *K_ic_* and effective length *l_ic_*. Parameter sets are chosen to reproduce the correlation–neighbor-order profiles observed experimentally in panel (A). Points and error bars indicate the mean ± SD. **(C)** Effects of inter-chromosomal spring effective length in simulations. Displacement correlation is plotted versus neighbor order for increasing values of spring effective length (*l_ic_*) at fixed spring stiffness (*K_ic_* = 0.5 pN *µ*m^−1^). Increasing effective length reduces the decay of correlations with neighbor order. Correlations are averaged over time lags from 5–50 s. Points and error bars indicate the mean ± SD. **(D)** Effects of inter-chromosomal spring stiffness in simulations. Displacement correlation is plotted versus neighbor order for increasing values of spring stiffness (*K_ic_*) at fixed effective length (*l_ic_* = 3 *µ*m). Changing stiffness primarily rescales correlation magnitude while preserving its dependence on neighbor order. Correlations are averaged over time lags from 5–50 s. Points and error bars indicate the mean ± SD. **(E)** Conceptual schematics summarizing how Noc and TSA reshape mechanical coupling between chromosomes. Left, untreated cells exhibit mechanical coupling mediated by the spindle together with relatively stiff, compact chromosomes, enabling efficient force propagation across the spindle. Middle, Noc-treated spindles are enriched in stabilized kinetochore microtubules and form a mechanically denser network, allowing forces to be transmitted more coherently across longer distances and extending coordination to higher neighbor orders. Right, TSA-treated cells exhibit softened, decondensed chromatin that locally dissipates mechanical stress, weakening elastic coupling between chromosomes. Subtle changes in microtubule mechanical properties may slow stress propagation, leading to delayed buildup of coordinated motion.

To identify which inter-chromosomal spring properties underlay these two effects, we systematically varied the spring effective length (*l_ic_*) and stiffness (*K_ic_*) in simulations. Increasing *l_ic_* primarily expanded the range of coordination, extending correlations across higher neighbor orders without strongly changing their peak amplitude (Figure 6C). In contrast, increasing *K_ic_* enhanced the magnitude of coordination without substantially altering its spatial decay (Figure 6D). The effect of *K_ic_*saturated at higher values, whereas increases in *l_ic_* continued to extend the spatial reach of coordination. Guided by these trends, we mapped simulation parameters that reproduced both the mean and variance of displacement correlations observed experimentally. Increasing *l_ic_*recapitulated the extended, long-ranged coordination induced by Noc treatment, whereas decreasing *K_ic_* reproduced the weakened coordination characteristic of TSA-treated cells (Figure 6B).

We next examined how coordination evolved over time by comparing displacement correlations as a function of time lag for first-order neighbors (Figure S4D). In Noc-treated cells, the temporal dependence of displacement correlations closely overlapped with control from 5 to 100 s. In contrast, TSA-treated cells exhibited a reduced plateau value and took longer to reach the plateau. Using model parameters calibrated to reproduce the experimentally observed coordination-versus-distance relationships, simulations successfully captured the reduced time-lagged correlation in the TSA condition but failed to reproduce the near-complete overlap between control and Noc curves (Figure S4E).

In all, this analysis framework allowed us to separate chromosome coordination into three key features: (i) how far correlations extend across the spindle, (ii) how strong coordination ultimately becomes, and (iii) how quickly coordination emerges over time. TSA treatment altered all three features of chromosome coordination, extending the spatial range of coupling while simultaneously reducing its strength and slowing its emergence. In contrast, Noc selectively increased the spatial extent of coordination without substantially affecting its magnitude or temporal evolution. In the discussion, we relate these three features to distinct mechanical properties of the spindle.

## Discussion

Chromosomes are dynamic throughout the cell cycle. Upon condensation and attachment to spindle microtubules in metaphase, individual chromosomes exhibit oscillatory motion. Oscillation of chromosome pairs is also coordinated, indicative of spatial coupling among chromosomes. We discovered that coordinated motion persists even when oscillation amplitude is substantially reduced. Our ability to separate oscillation strength from coordinated motion suggests that coordination does not arise from oscillations themselves, but from forces between individual chromosome pairs. Through a minimal mechanical model, we demonstrate that transient, stochastic inter-chromosomal connections are sufficient to generate coordinated motion, revealing that coordination can emerge purely from shared mechanical linkages. Notably, perturbations to microtubules and chromatin produced distinct changes in the spatial pattern of coordination, indicating that these two systems support inter-chromosomal coupling through different physical mechanisms. In the sections below, we discuss experimental observations and modeling results to discern how microtubule-based and chromatin-based pathways mechanistically shape both the oscillatory dynamics of individual chromosomes and the spatial organization of coordination across the metaphase plate.

### Minimal Model Reveals Mechanical vs. Regulatory Contributions to Chromosome Dynamics

We developed a minimal mechanical model to isolate the extent to which chromosome dynamics can emerge from force balance alone, independent of detailed molecular regulation. In the model, chromosome motion is governed by elastic coupling between sister kinetochores through the centromere, which provides a restoring force that opposes the poleward pulling forces generated at each kinetochore. Rather than explicitly modeling individual microtubule growth and shortening events, we represent kinetochore force generation using a tension-dependent effective number of kinetochore–microtubule attachments. This attachment number reacted to chromosome motion and centromere stretch, establishing a feedback loop in which kinetochore force strengthens during poleward motion, until stretch of the centromeric spring ultimately reverses motion, generating sustained oscillations. This abstraction preserves the essential mechanical feedback underlying chromosome oscillations while substantially reducing model complexity. Prior theoretical work has shown that oscillatory chromosome motion can arise from force–balance feedback alone (Banigan et al., 2015; Schwietert and Kierfeld, 2020). In particular, Banigan et al. (2015) demonstrated that each kinetochore behaves as a bistable force generator, producing weak pulling forces and moving slowly in the anti-poleward direction under low load, while switching to a strong pulling state that drives rapid poleward motion under high load. The load on each kinetochore is set by centromere stretch. Similar to our model, oscillations arise from feedback between load-dependent force generation at the kinetochore and elastic coupling via the centromere: when both sisters move in the same direction, centromere stretch is reduced, destabilizing the force state at the leading kinetochore and triggering a reversal in motion. Our model captures this same mechanical feedback without explicitly resolving microtubule growth and shortening states, thereby distilling the framework into a more coarse-grained form.

A key advantage of a minimal model is that, by reducing the system to its mechanical essentials, it distinguishes features of chromosome behavior that arise through force balance from those that require additional molecular regulation. Our model substantially underestimates the amplitude and variability of centromere stretch compared with experiment (Figure 2C), indicating that centromere stretch dynamics are shaped by regulatory mechanisms not captured in a purely mechanical description. Prior work in PTK1 cells has shown that kinetochore motion is intrinsically asymmetric, with poleward movement occurring faster than anti-poleward movement. This velocity asymmetry causes the two sister kinetochores to switch direction at different times, creating a transient window of differential motion during which centromere stretch accumulates (Wan et al., 2012). Substantial centromere extension therefore requires a sufficiently prolonged time lag between sister kinetochore motions, such that one kinetochore advances poleward while the other remains delayed. In our minimal model, centromere stretch likewise arises from differential sister motion, but these differences are short-lived. Because kinetochore–microtubule attachment number changes proportionally and symmetrically between sister kinetochores in response to velocity, load is redistributed rapidly once centromere tension builds, producing only a brief time lag and tightly constraining centromere extension. Together, these results suggest that while oscillatory switching itself is mechanically driven, large and variable KK oscillation amplitudes require additional layers of regulation that prolong the duration of differential motion between sister kinetochores.

### Relationship between chromosome oscillations and coordination

Chromosome coordination refers to the correlated movement of spatially separated chromosomes within the metaphase spindle, such that their motions are linked in time and direction. A key experimental finding of this study is that chromosome coordination persists even when oscillatory amplitude is strongly dampened by perturbations to either microtubules or chromatin. Low-dose nocodazole and histone hyperacetylation both suppress regular oscillatory behavior while having no effect on coordinated motion between chromosomes. This observation supports a model in which chromosome coordination reflects mechanical coupling through the spindle environment, allowing motion to be transmitted between chromosomes even when oscillations are weak or stochastic. Consistent with this interpretation, our simulations show that inter-chromosomal coordination is largely insensitive to the parameters that regulate individual chromosome activity, including centromere stiffness and KT–MT turnover rates (Figure S3B). Together, these findings indicate that oscillatory motion and coordination are governed by distinct mechanical principles: oscillations reflect local force balance at individual kinetochores, whereas coordination reflects how forces propagate through the surrounding spindle.

### Inter-chromosomal springs as an effective description of spindle-mediated force transmission

To investigate mechanical coupling mechanisms that could give rise to coordinated chromosome motion, we incorporated stochastic springs between neighboring chromosomes as a coarse-grained representation of force transmission through the spindle medium, rather than direct chromosome–chromosome attachments. While previous studies have reported coordinated metaphase chromosome movements and implicated specific spindle-associated proteins, including motor proteins, in mediating this behavior (Vladimirou et al., 2013), our results show that coordination does not require dedicated molecular crosslinkers between chromosomes. Instead, coordinated motion emerges naturally when chromosomes are mechanically coupled through a shared physical environment. Within this framework, both the magnitude and structure of coordination are governed by the dynamics and physical properties of the coupling itself (Figure 4C). To relate chromosome motion quantitatively to the mechanical properties of this coupling, we adopted a two-point microrheology–inspired analysis. In this interpretation, inter-chromosomal springs encode effective viscoelastic properties of the spindle. The spring length defines a characteristic length scale that sets how far stresses propagate through the spindle, thereby controlling the spatial range of coordination. Spring stiffness reflects the effective elastic response of the system to relative chromosome displacements, tuning the amplitude of correlated motion without substantially altering its spatial range (Figure 6C–D). From this systems-level perspective, chromosome coordination emerges from the collective mechanics of the spindle material itself. Consistent with this view, work from the Needleman group has demonstrated that chromosomes embedded within mammalian spindles can exhibit long-range spatial anti-correlations, reflecting repulsive mechanical interactions mediated by the spindle material itself rather than direct molecular linkages between chromosomes (Kelleher et al., 2024).

Guided by these principles, we interpret the distinct effects of nocodazole (Noc) and trichostatin A (TSA) on chromosome coordination in terms of how each perturbation reshapes spindle-mediated force transmission (Figure 6E). Following Noc treatment, the spindle becomes more compact and enriched in stabilized microtubules (Figure S1C), generating a denser environment in which forces are transmitted more continuously across chromosomes. Consistent with this interpretation, coordination extends over larger neighbor orders after Noc treatment (Figure 6A). In contrast, TSA induces pronounced chromatin decondensation, rendering chromosomes softer and more compliant (Spicer and Gerlich, 2023; Biggs et al., 2019). In this softened regime, inter-chromosomal mechanical interactions are less effective at transmitting force, as local deformation dissipates stress rather than propagating it through the spindle network. Accordingly, TSA reduces the overall magnitude of coordination, consistent with weakened elastic coupling between chromosomes (Figure 6A). Notably, TSA also uniquely alters the temporal buildup of coordination: displacement correlations take longer to reach a plateau following TSA treatment (Figure S4D), indicating slower stress propagation and increased viscous dissipation within the spindle environment. This delayed temporal response is likely linked to TSA’s secondary effects on microtubule mechanics through inhibition of HDAC6 and the resulting increase in *α*-tubulin acetylation. Acetylated microtubules are longer-lived and less dynamic, and HDAC6 inhibition has been shown to suppress microtubule turnover (Perdiz et al., 2011; Asthana et al., 2013; Li and Yang, 2015).

While both Noc and TSA are expected to stabilize microtubules, Noc treatment leads to an increase in spindle density that is not observed following TSA treatment. Instead, TSA is likely to alter microtubule mechanical properties. Previous studies have shown that elevated *α*-tubulin acetylation increases microtubule compliance, resilience to mechanical stress, and internal dissipation (Xu et al., 2017; Portran et al., 2017). Under TSA treatment, a more compliant and dissipative microtubule network would be expected to slow stress propagation through the spindle while allowing correlations to extend over longer spatial ranges. In contrast, following Noc treatment, increased microtubule density promotes more continuous force transmission, extending the spatial range of chromosome coordination without substantially altering its temporal dynamics.

Together, our results identify chromosome coordination as an emergent mechanical property of the mitotic spindle rather than a consequence of synchronized oscillatory programs or dedicated chromosome–chromosome linkers. By decoupling oscillatory dynamics from coordination, we show that individual chromosome motion is governed by local force balance at kinetochores, whereas coordination reflects force transmission through the spindle as a mechanically coupled, viscoelastic material. Within this framework, microtubules primarily set the spatial continuity of force transmission and the temporal buildup of coordination, while chromatin mechanics tune the strength of coordination. Coordinated chromosome motion therefore serves as a systems-level readout of spindle material properties, integrating contributions from multiple structural components rather than any single molecular pathway. We propose that this distributed mechanical coupling stabilizes chromosome positioning during metaphase by enabling long-range force integration that dampens stochastic fluctuations and preserves global spindle organization.

## Materials and Methods

### Cell Culture and Drug Treatments

PTK1 cells stably expressing Hec1–GFP were originally obtained from the DeLuca laboratory (DeLuca et al., 2018). Cells were maintained at 37 ^◦^C and 5% CO_2_ in a humidified incubator and cultured in DMEM/F12 medium (Thermo Fisher Scientific) supplemented with 10% fetal bovine serum (FBS; Thermo Fisher Scientific) and 100 U/mL penicillin–streptomycin (Life Technologies). Cells were passaged every 3–4 days and used for experiments at 80–90% confluence.

For drug perturbations, cells were treated with nocodazole (10 ng/mL; Cell Signaling Technology) for 1 h or trichostatin A (500 nM; Sigma) for 3 h prior to live-cell imaging or fixation. For both live-cell and fixed-cell imaging experiments, cells were plated on 4-well glass-bottom dishes with high-performance #1.5 cover glass (CellVis) three days prior to imaging or fixation.

### Live-Cell Imaging

Live-cell time-lapse imaging of PTK1:Hec1–GFP cells for kinetochore tracking was performed on a custom-built light-sheet fluorescence microscope as previously described (Fadero et al., 2018). Images were acquired using a Nikon Plan Apochromat 100×/1.35 NA silicone immersion objective, a Photometrics Prime 95B sCMOS camera, and a stage-top incubator (Tokai Hit).

Image volumes were collected with a z-step size of 0.5 *µ*m over a total axial range of 3 *µ*m, with one volume acquired every 5 s. Imaging was initiated in early metaphase, when most sister kinetochore pairs were aligned at the metaphase plate, and continued until anaphase onset. For all cells, an ROI of identical size was selected around the mitotic cell to ensure uniform spatial resolution across imaging datasets.

### Quantification of chromosome, Spindle, and Acetylated Histone Fluorescence

PTK1 cells were fixed with 4% paraformaldehyde (Electron Microscopy Sciences) and processed for immunofluorescence using standard procedures, including permeabilization with 0.1% Tween-20 in PBS, and blocking with 1% bovine serum albumin (BSA, Sigma) in PBS. Acetylated histones were detected using an anti–acetyl-Histone H3 antibody (Sigma), chromosomes were labeled with DAPI (Sigma), and spindle microtubules were visualized by expression of Halo-tagged tubulin, which was fluorescently labeled using a Halo-dye ligand (Halo-JFX650, gift from Janelia).

Three-dimensional fluorescence image stacks encompassing the full mitotic volume were acquired using identical imaging parameters (illumination, exposure time, detector gain, pixel size, and z-step) for all experimental conditions. Chromosome, spindle, and acetylated histone fluorescence signals were quantified from 3D image stacks using custom ImageJ/Fiji macros. Chromosome volumes were segmented from the DAPI signal, spindle regions from the microtubule channel, and acetylated histone signal from the corresponding antibody channel. For each segmented region of interest, the total integrated fluorescence intensity and the number of voxels were measured, and fluorescence intensity was reported as mean intensity per voxel (A.U./voxel) to normalize differences in segmented volume. Background intensity was measured from cell-free regions and subtracted prior to quantification. All analyses were performed using identical thresholds and analysis parameters across conditions within each experiment.

### Kinetochore Tracking

Individual sister kinetochore positions were tracked over time from Hec1–GFP fluorescence using TrackMate (ImageJ/Fiji). Prior to tracking, image stacks were rotated such that the two spindle poles were aligned horizontally, allowing kinetochore motion to be quantified along a single axis (x-direction). Kinetochore foci were detected as diffraction-limited spots using a Gaussian detector with an estimated spot diameter of 0.5 *µ*m. Spots were linked between successive frames using the TrackMate linear assignment tracker with a maximum linking distance of 1 *µ*m. Tracks were manually inspected to correct occasional linking errors, and only kinetochore tracks present for the full imaging duration were included in subsequent analysis.

Sister kinetochore pairs were assigned manually by identifying pairs of tracks that remained in close spatial proximity and exhibited highly correlated motion over time, consistent with known sister kinetochore behavior.

### Quantification of Chromosome Oscillation and Activity

To correct for global spindle movement, kinetochore positions were expressed in a pole-aligned coordinate system, with one spindle pole defined as the reference origin (*x* = 0). Individual kinetochores were tracked and used as proxies for chromosome motion. Each chromosome pair contains two sister kinetochores, which were tracked independently. Chromosome-pair motion was represented by two derived trajectories: the center-of-mass (CM) trajectory, defined at each time point as the mean of the pole-aligned x-positions of the two sister kinetochores, and the inter-kinetochore (KK) distance, defined as the difference between the two kinetochore positions.

Chromosome oscillatory behavior was quantified from both CM and KK trajectories by identifying local maxima and minima over time. Oscillation period was calculated as the time interval between successive CM peaks (or valleys), and oscillation amplitude was quantified from peak-to-valley excursions within individual oscillation cycles. Measurements were computed separately for CM and KK oscillations for each chromosome pair and averaged across oscillation cycles to obtain a single period and amplitude for each metric.

Under experimental conditions in which chromosome oscillations were substantially dampened, overall chromosome activity was quantified as an alternative metric of chromosome dynamics. For this analysis, instantaneous kinetochore velocities were computed from pole-aligned kinetochore trajectories. For each sister kinetochore trajectory 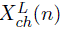 and 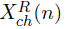, instantaneous velocities were estimated using a sliding three-frame linear fit. At each time index *n*, a least-squares line was fit to the three consecutive position measurements {(*t_n_, X*(*n*)), (*t_n_*_+1_*, X*(*n* + 1)), (*t_n_*_+2_*, X*(*n* + 2))}, and the slope of this fit was taken as the instantaneous velocity at time *t_n_*. Velocities were computed separately for the left and right kinetochores and aligned by time. Chromosome-pair activity was defined as the time-averaged sum of the squared kinetochore velocities,

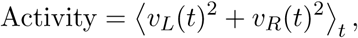

with units of *µ*m^2^ min^−2^.

### Quantification of Chromosome Coordination

Chromosome coordination was quantified using correlations between chromosome-pair center-of-mass (CM) trajectories. Coordination was quantified using a normalized cross-correlation function. For two CM trajectories *x*_1_(*t*) and *x*_2_(*t*), the cross-correlation at time lag *τ* was computed as

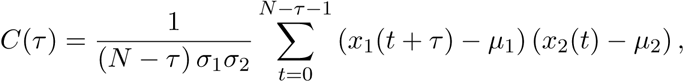

where *µ*_1_, *µ*_2_ and *σ*_1_, *σ*_2_ are the means and standard deviations of the two trajectories, and N is the number of time points in the trajectory.

Only trajectories with at least 20 frames were analyzed. The zero-lag value of this normalized cross-correlation, *C*(0), was used as the coordination metric. Because the trajectories are mean-subtracted and variance-normalized, *C*(0) is equivalent to the Pearson’s correlation coefficient (R) between the two CM trajectories. Correlations were computed for all possible chromosome-pair combinations within each spindle, and results were pooled across spindles for population-level analysis.

### Single-Chromosome Mechanical Model

To investigate the minimal mechanical requirements for chromosome oscillatory motion, we developed a one-dimensional, coarse-grained mechanical model of a single metaphase chromosome pair consisting of two sister chromatids. Each chromatid is represented by a point mass corresponding to its kinetochore-associated chromosome position, denoted 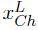 and 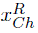. Motion occurs along the spindle axis only; transverse motion is neglected.

<u>Overdamped Force Balance</u> Chromosome motion is assumed to be overdamped and governed by a balance between viscous drag, kinetochore–microtubule (KT–MT) pulling forces, and centromeric restoring force. The equations of motion for the left (L) and right (R) chromatids are given by

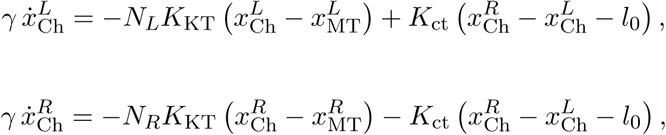

where 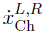 are chromatid velocities, *γ* is the effective viscous drag coefficient, *K*_KT_ is the effective stiffness of individual KT–MT attachments, *N_L,R_* are the numbers of KT-MT attachments on each kinetochore, *K*_ct_ is the centromeric spring stiffness, and *l*_0_ is the centromere rest length.

<u>Kinetochore–Microtubule Attachment Dynamics</u> The number of KT–MT attachments evolves dynamically in response to chromosome motion. Attachments form with a baseline rate *n*, increase with load-dependent stabilization, saturate at a maximum attachment number *N*_max_, and detach with rate *λ*. The attachment dynamics are modeled as

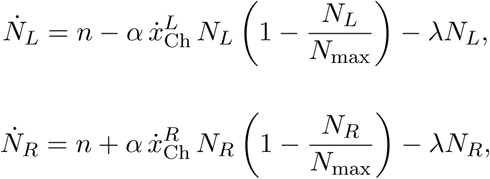

where *α* sets the coupling between chromosome motion and attachment stabilization. Faster poleward motion increases load on kinetochore–microtubule linkages at the leading kinetochore, stabilizing additional attachments and amplifying pulling forces until centromeric restoring forces dominate and reverse motion, closing the feedback loop.

Microtubule tip motion The positions of the microtubule plus-ends at the kinetochores, 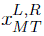, were assumed to move proportionally to chromosome velocity, representing collective microtubule growth and shortening:

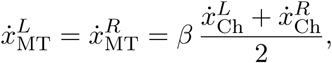

where *β* is a scaling factor controlling the coupling between chromosome motion and microtubule tip motion.

<u>Stochasticity and numerical integration</u> To capture variability in chromosome motion, stochastic fluctuations were incorporated in two ways. First, small multiplicative noise was added to selected kinetic parameters (*n*, *α*, *λ*) at the start of each simulation to represent cell-to-cell variability. Second, additive Brownian noise was applied to chromosome positions at each time step to produce diffusive jitter that contributes to both center-of-mass (CM) and inter-kinetochore (KK) fluctuations. The coupled equations were integrated numerically using a forward Euler scheme with time step *dt*. Simulations were performed for *N*_steps_ iterations, and trajectories of chromatid positions, attachment numbers, center-of-mass position,

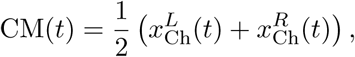

and inter-kinetochore distance

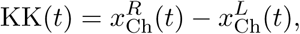

were recorded.

<u>Model outputs and analysis</u> Simulated CM and KK trajectories were analyzed using the same oscillation and activity quantification procedures applied to experimental data. Oscillation amplitude and period were extracted from CM and KK time series using peak– and valley-based analysis, and chromosome activity was quantified from kinetochore velocities by computing the time-averaged sum of squared instantaneous velocities of the two sister kinetochores. Simulation results were exported for downstream analysis and direct comparison with experimental measurements.

### Inter-Chromosomal Coupling Model

To investigate how mechanical interactions between chromosomes generate coordinated motion, the single-chromosome model was extended to simulate *N_chromosome_*pairs embedded in a shared spindle environment. Each chromosome pair *i* consists of two sister chromatids with positions 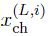 and 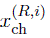, whose dynamics follow the same overdamped force balance and kinetochore–microtubule attachment kinetics described in the single-chromosome model. Motion was restricted to one dimension along the spindle axis, while chromosome pairs were assigned fixed, evenly spaced positions along the transverse axis for visualization purposes only.

Inter-chromosomal mechanical coupling

In addition to forces acting within individual chromosome pairs, chromosomes were coupled to one another through transient, pairwise mechanical connections that transmit forces between chromosome centers of mass. For chromosome pair *i*, the equations of motion become

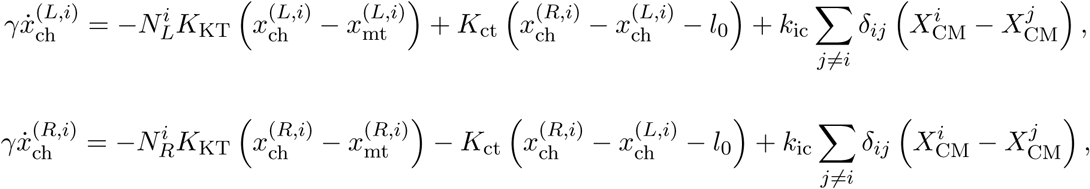

where *k*_ic_ is the inter-chromosomal coupling stiffness, *δ_ij_* ∈ {0, 1} denotes the instantaneous presence or absence of a mechanical connection between chromosome pairs *i* and *j*. 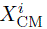 and 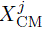 are the center-of-mass positions of chromosome pairs *i* and *j*, respectively.

<u>Stochastic inter-chromosomal connectivity</u> Mechanical coupling between chromosomes was mediated by transient, stochastic connections formed between the centers of mass (CM) of chromosome pairs. For each chromosome pair (*i, j*), the instantaneous inter-chromosomal distance was computed as

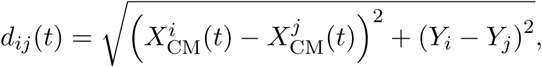

where *X*_CM_ denotes the center-of-mass position along the spindle axis and *Y* is the fixed transverse position assigned to the chromosome.

The connection state between chromosomes *i* and *j* was represented by a binary variable *δ_ij_*(*t*) ∈ {0, 1}, indicating whether a mechanical connection was present. Inter-chromosomal connections formed and dissociated stochastically with rates that depended on the inter-chromosomal distance *d_ij_*. Specifically, the distance-dependent association and dissociation rates were defined as

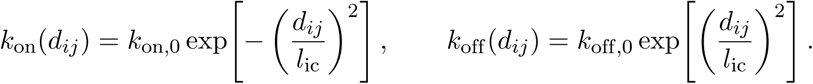

Here, *k*_on,0_ and *k*_off,0_ are the base association and dissociation rates, respectively, which set the maximal probabilities of connection formation and breakage at short inter-chromosomal distances. The parameter *l*_ic_ defines the effective interaction length of inter-chromosomal springs, setting the spatial range over which chromosomes can connect. Once connected, springs have zero mechanical rest length, such that the coupling force is zero only when 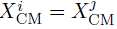 and increases with inter-chromosomal separation. Consequently, the coupling does not impose a preferred spacing but solely resists relative motion between chromosomes. At each simulation time step *dt*, new connections formed with probability *k*_on_(*d_ij_*) *dt*, while existing connections dissociated with probability *k*_off_ (*d_ij_*) *dt*. Formation and dissociation events were treated as mutually exclusive, yielding a dynamically evolving adjacency matrix *δ_ij_*(*t*) that captured the stochastic network of inter-chromosomal connections over time.

<u>Stochasticity and numerical integration</u> To represent chromosome-to-chromosome variability within a spindle, kinetochore–microtubule attachment parameters (*n_i_, α_i_, λ_i_*) were drawn independently for each chromosome pair *i* at the start of each simulation, using the same multiplicative noise model as in the single-chromosome case. Additive Brownian noise was applied to chromosome positions at each time step, producing diffusive jitter that contributed to both center-of-mass (CM) and inter-kinetochore (KK) fluctuations for each chromosome. The coupled system of differential equations for all chromosomes, including inter-chromosomal coupling forces, was integrated numerically using a forward Euler scheme with time step *dt*. Simulations were run for *N*_steps_ iterations, yielding time-resolved trajectories of left and right chromatid positions for each chromosome, attachment numbers, and evolving inter-chromosomal connectivity matrix *δ_ij_*(*t*).

Model output and analysis For each chromosome pair *i*, center-of-mass and inter-kinetochore coordinates were computed as

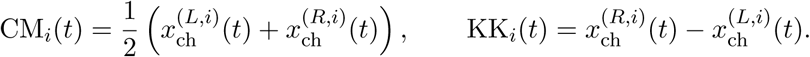

Simulated CM and KK trajectories for each chromosome were analyzed using the same oscillation and activity quantification procedures applied to both the single-chromosome model and experimental data. In addition to characterizing per-chromosome dynamics, the multi-chromosome model enabled direct quantification of chromosome coordination. Coordination was assessed using two complementary approaches: (i) Pearson correlation analysis of chromosome-pair CM trajectories, as described in the Chromosome coordination quantification section, and (ii) time-lagged displacement correlation analysis, described below. Correlation measurements were computed for all chromosome pairs within a spindle and pooled across simulation replicates to obtain ensemble coordination statistics.

### Time-Lagged Displacement Correlation Analysis

To quantify chromosome coordination across defined temporal and spatial scales, we implemented a time-lagged displacement correlation analysis inspired by two-point microrheology. This approach complements Pearson’s R measurements by resolving coordination as a function of both time lag and spatial separation, while increasing statistical power by pooling displacement measurements across time windows and chromosome pairs.

For each cell, the center-of-mass (CM) position of every chromosome pair was computed from sister-kinetochore trajectories as described above. Let *X_i_*(*t*) denote the CM position of chromosome *i* at time *t*. For a given time lag *τ*, time-lagged displacements were calculated as

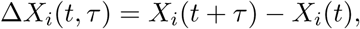

with *t* indexing all time points for which the lagged displacement is defined. Each data point corresponds to a 5 s imaging interval; lag times from *τ* = 1 to *τ* = 30 therefore correspond to physical times of 5–150 s.

For each unique chromosome pair (*i, j*) with *i* ≠ *j*, displacement coordination at lag *τ* was quantified as the time-averaged product of their displacements,

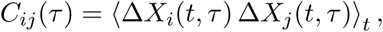

where the average was taken over all valid time points within a trajectory.

To relate coordination to spatial separation while minimizing noise from instantaneous distance fluctuations between chromosomes, chromosome pairs were grouped by neighbor order, defined by their relative ordering along the metaphase plate. First-order neighbors correspond to immediately adjacent chromosomes along the metaphase plate, with higher neighbor orders indicating progressively larger separations. For each lag *τ* and neighbor order *n*, displacement correlations were averaged over all chromosome pairs belonging to that neighbor order and over all time points,

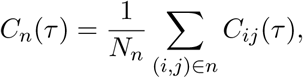

where *N_n_* denotes the number of chromosome pairs in neighbor order *n*. Correlation values were computed for each cell individually and then pooled across cells to obtain population-averaged coordination statistics. For each neighbor order *n* and time lag *τ*, data are reported as the mean ± standard deviation across cells.

### Statistical Analysis

Statistical analyses were performed in Python using SciPy and Statsmodels. Data were analyzed using one-way analysis of variance (ANOVA) when comparing metrics across multiple experimental conditions, including live-cell chromosome activity measurements and fixed-cell measurements, such as histone acetylation signal intensity, spindle microtubule area and fluorescence intensity, and chromosome area and fluorescence intensity. When a significant overall effect was detected by ANOVA, post hoc multiple comparisons were performed using Tukey’s honestly significant difference (HSD) test to correct for multiple comparisons. All tests were two-sided, and a significance level of 0.05 was used.

Unless otherwise stated, data points represent individual cells or chromosome pairs, as appropriate for each analysis. Statistical significance is reported as *, *P <* 0.05; **, *P <* 0.01; ***, *P <* 0.001; and ns, not significant.

## Acknowledgments

We are grateful to the DeLuca laboratory at Colorado State University for generously providing the PTK1:Hec1–GFP cell line. This work was supported by the U.S. Department of Energy under grant A23-1683-001.

## Author contributions

J.Z. conceived the study, designed and performed experiments, developed and implemented all computational models and analysis pipelines, analyzed the data, and wrote the manuscript.

## Data and code availability

All simulation and analysis code supporting this study is publicly available at https://github.com/Jiali-Z/Coordinated_Chromosome_Oscillation (release v1.0.0).

