## Supplemental Information for "Coordinated chromosome motion emerges from mechanical coupling mediated by the physical spindle environment"

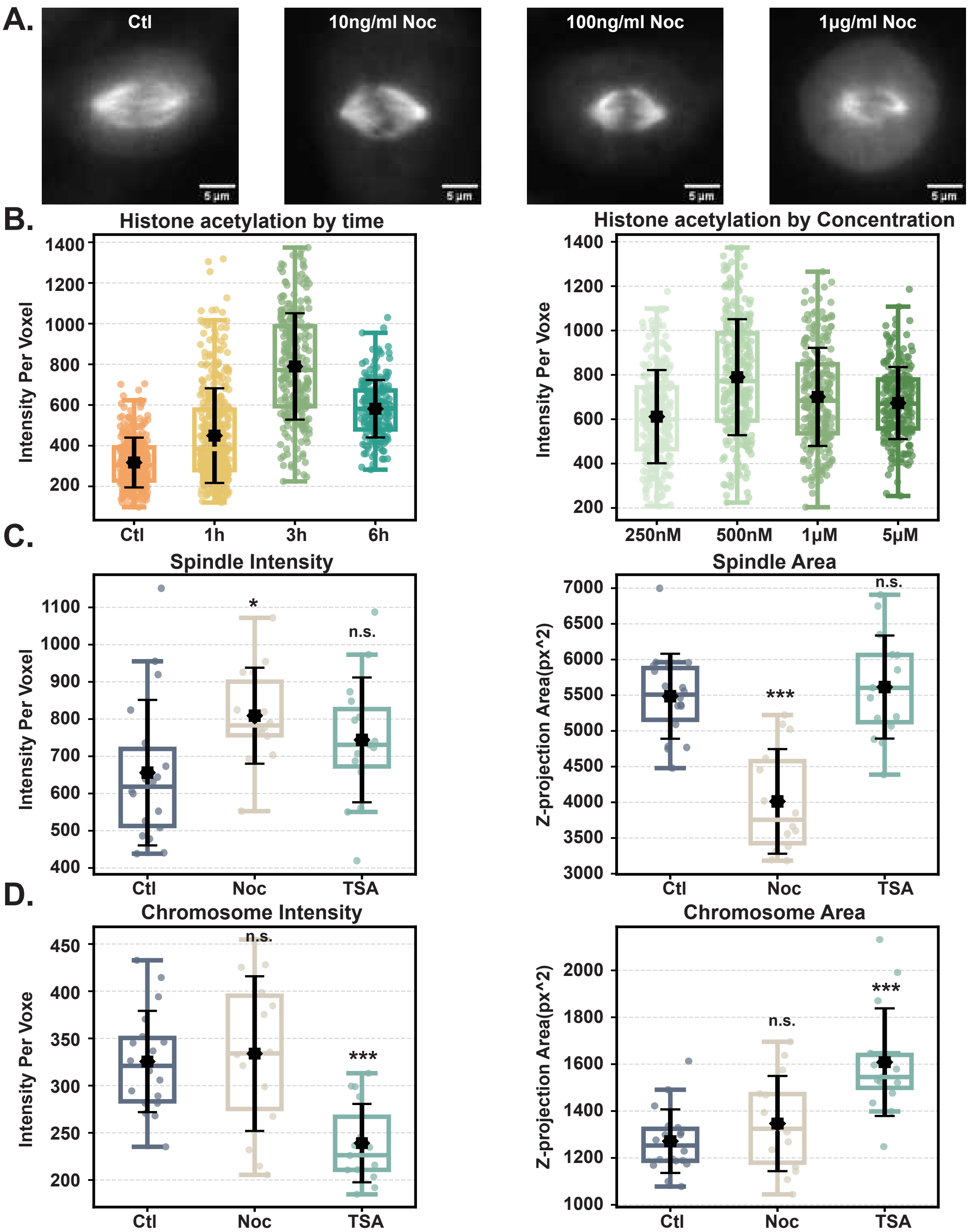


**Figure S1. Low-dose nocodazole alters spindle architecture, whereas trichostatin A induces chromatin decondensation.**

**Figure S1. Low-dose nocodazole alters spindle architecture, whereas trichostatin A induces chromatin decondensation. (A)** Representative live-cell images of metaphase PTK1 cells treated with nocodazole (Noc) at the indicated concentrations ($1\ \mu g\,mL^{-1}$,$100\ ng\,mL^{-1}$,$10\ ng\,mL^{-1}$), and untreated control (Ctl). Images show spindle morphology under each condition. Scale bars, 5 $\mu$m. **(B)** Quantification of histone acetylation levels as a function of trichostatin A (TSA) treatment time (left) and TSA concentration (right), measured as mean voxel intensity. Treatment with 500 nM TSA for 3 h yielded the largest increase in histone acetylation and was used for subsequent experiments. **(C)** Quantification of spindle properties under control (Ctl), Noc-treated ($10\ ng\,mL^{-1}$, 1 h), and TSA-treated (500 nM, 3 h) conditions. Spindle microtubules were visualized in fixed PTK1 cells expressing Halo-tagged tubulin and fluorescently labeled with 10 nM Halo-JFX dye for 5 min at 37 °. Left, spindle microtubule intensity increases following Noc treatment, whereas TSA produces no significant change. Right, nocodazole significantly reduces spindle area, while TSA does not significantly alter spindle size. **(D)** Quantification of chromosome properties under control, Noc, and TSA conditions. Chromosomes were visualized by treating in fixed PTK1 cells by staining with DAPI (1:1000) for 10 min at room temperature. Left, chromosome intensity is significantly reduced after TSA treatment but unchanged following Noc. Right, TSA induces an increase in chromosome area, whereas Noc does not significantly affect chromosome area. Each point represents one cell. Boxplots show the distribution, with black symbols indicating the mean ± SD. Statistical significance is indicated as shown ($^{*}p < 0.05$; $^{***}p < 0.001$; n.s., not significant).

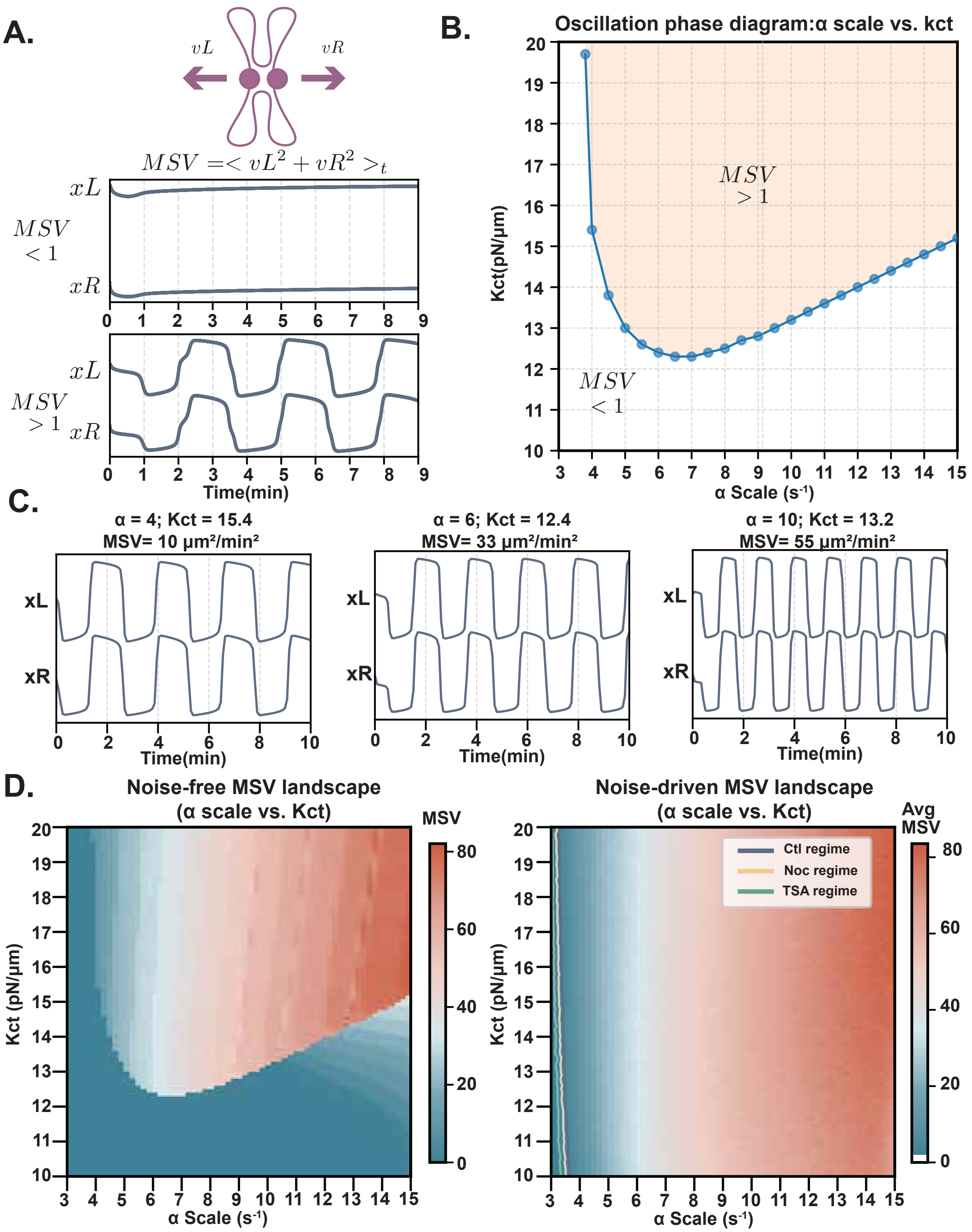


**Figure S2. Deterministic and noise-driven regimes of chromosome oscillations in the minimal single-chromosome mechanical model.**

**Figure S2. Deterministic and noise-driven regimes of chromosome oscillations in the minimal single-chromosome mechanical model. (A)** Schematic and representative trajectories illustrating how chromosome activity is classified in the model using the mean squared velocity (MSV). Left and right kinetochore positions ($x_L$ and $x_R$) are shown for parameter regimes that produce non-oscillatory motion (MSV $< 1$; top) or sustained oscillations (MSV $> 1$; bottom) in the deterministic version of the model. MSV serves as a scalar measure of overall chromosome activity, with larger values indicating stronger and more persistent oscillatory motion. **(B)** Oscillation phase diagram showing regions of parameter space that support oscillatory behavior in the deterministic model, plotted as centromere spring stiffness ($K_{\mathrm{ct}}$) versus the kinetochore-microtubule attachment dynamics ($\alpha$). The boundary separates non-oscillatory (MSV $< 1$) and oscillatory (MSV $> 1$) regimes. At lower values of $\alpha$, oscillations require weaker centromere springs, whereas increasing $\alpha$ shifts the oscillatory regime toward larger $K_{\mathrm{ct}}$ values. **(C)** Representative deterministic trajectories from three locations within the oscillatory regime of the phase diagram. As $\alpha$ and $K_{\mathrm{ct}}$ increase, oscillations become faster and more frequent, while their overall amplitude changes only modestly. Parameter values and corresponding MSV values are indicated above each trace. **(D)** Simulated MSV landscapes across $\alpha$ and $K_{\mathrm{ct}}$ for the deterministic (noise-free; left) and noise-driven (right) versions of the model. In the absence of noise, a sharp boundary separates low- and high-activity regimes. When noises are included, this transition is smoothed, producing a graded increase in MSV across parameter space. Contours indicate regions corresponding to control (Ctl), nocodazole-treated (Noc), and trichostatin A–treated (TSA) regimes, which cluster within a similar range of $\alpha$ values ($\alpha = 3$–$4$ s$^{-1}$).

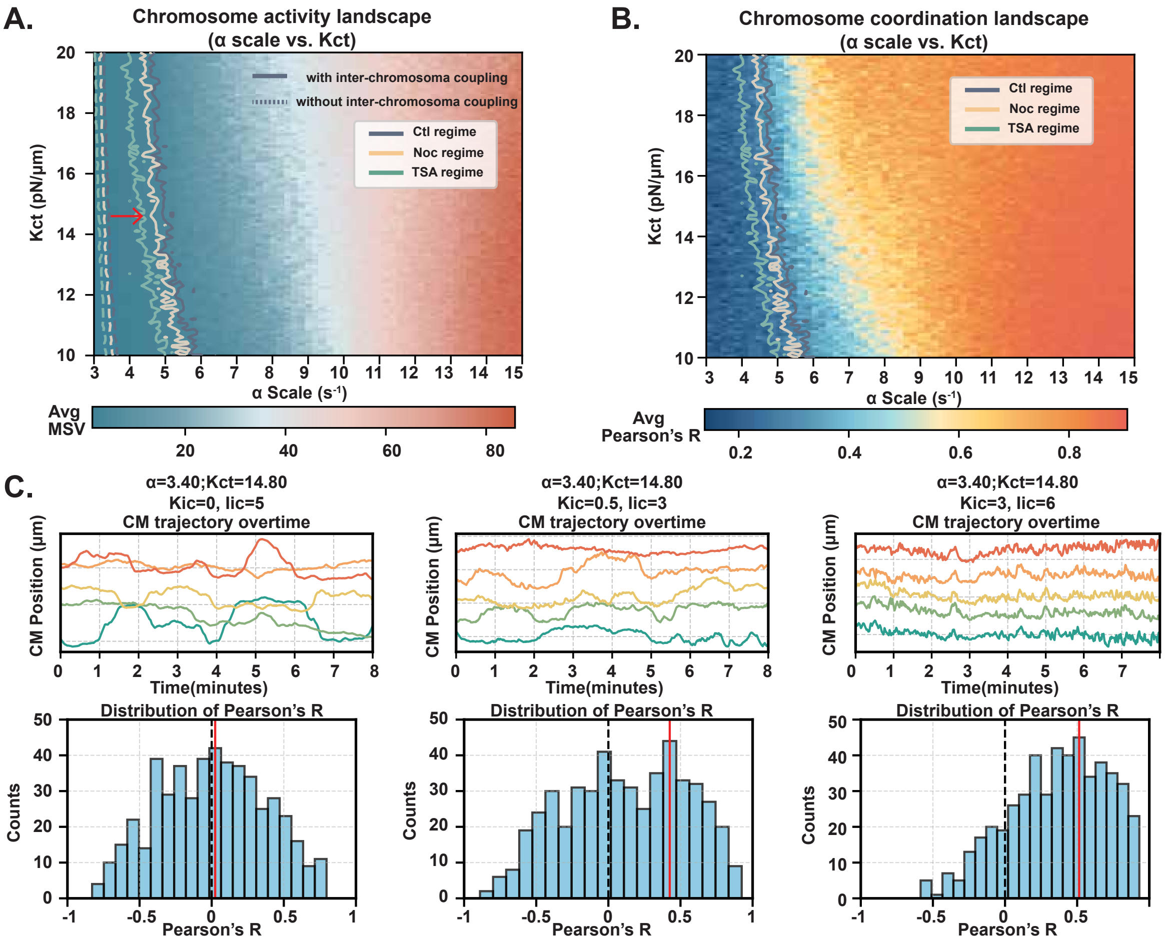


**Figure S3. Inter-chromosomal coupling alters chromosome coordination without strongly modifying single-chromosome activity.**

**Figure S3. Inter-chromosomal coupling alters chromosome coordination without strongly modifying single-chromosome activity. (A)** Chromosome activity landscape across the parameter space of centromere spring stiffness ($K_{\mathrm{ct}}$) and kinetochore-microtubule attachment rate ($\alpha$) in the presence of inter-chromosomal springs. The heat map shows the average mean squared velocity (MSV). Solid contours indicate parameter combinations corresponding to the experimentally defined control (Ctl), nocodazole-treated (Noc), and trichostatin A–treated (TSA) activity regimes. For comparison, dashed contours show the corresponding activity boundaries obtained from simulations without inter-chromosomal coupling (reproduced from Fig. S2D). Inter-chromosomal coupling shifts MSV contours upward (toward higher $\alpha$), indicating that stronger kinetochore-microtubule dynamics are required to achieve the same chromosome activity level when chromosomes are mechanically coupled. **(B)** Chromosome coordination landscape across the same kinetochore-microtubule attachment rate ($\alpha$) and centromere stiffness ($K_{\mathrm{ct}}$) parameter space. The heat map shows the average Pearson's R between chromosome center-of-mass trajectories. Overlaid contours correspond to the experimentally defined chromosome activity (MSV) regimes identified in panel (A) and are shown here for comparison. Pearson's correlation remains relatively uniform along these MSV contours, indicating that chromosome coordination is comparatively insensitive to variations in $\alpha$ and $K_{\mathrm{ct}}$ over the range of parameters that reproduce experimentally observed activity levels. **(C)** Effect of inter-chromosomal coupling on individual chromosome dynamics at fixed activity. Simulations were performed at $\alpha$ and $K_{\mathrm{ct}}$ values that reproduce experimentally measured chromosome activity in the single-chromosome model ($\alpha = 3.40\ \mathrm{s}^{-1}$, $K_{\mathrm{ct}} = 14.8\ \mathrm{pN}\,\mu\mathrm{m}^{-1}$). Inter-chromosomal coupling strength was varied by changing spring stiffness and resting length, as indicated. Top, representative center-of-mass (CM) trajectories of individual chromosomes over time. Bottom, corresponding distributions of Pearson's correlation coefficients pooled across simulation runs. Inclusion of inter-chromosomal springs enhances stochastic, diffusive fluctuations in individual chromosome motion and reduces the coherence of oscillatory trajectories, despite maintaining comparable overall chromosome coordination levels.

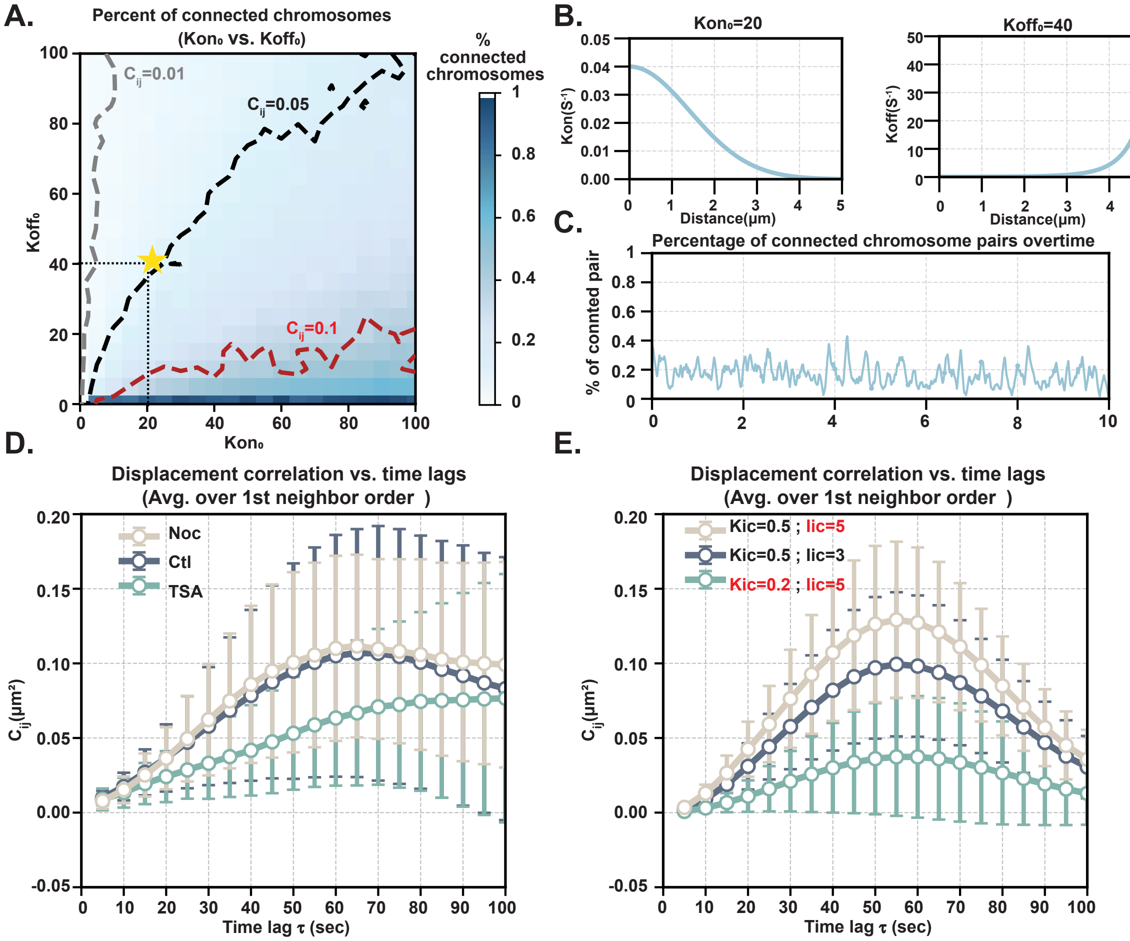


**Figure S4. Calibration of inter-chromosomal spring dynamics and displacement correlations versus time lag.**

**Figure S4. Calibration of inter-chromosomal spring dynamics and displacement correlations versus time lag. (A)** Percentage of chromosome pairs connected by inter-chromosomal springs across the space of attachment ($k_{\text{on}}$) and detachment ($k_{\text{off}}$) rates. The heat map shows the time-averaged fraction of connected chromosome pairs. Overlaid contours indicate values of the first-neighbor displacement correlation measured at the shortest time lag (5-50 s), with contours corresponding to $C_{ij} = 0.01$, 0.05, and 0.1. The yellow star marks the representative parameter combination ($k_{\text{on}} = 20$, $k_{\text{off}} = 40$) selected for subsequent simulations, which reproduces the experimentally observed control-level coordination. **(B)** Distance dependence of inter-chromosomal spring dynamics for the selected parameter set. Left, the attachment rate decreases with increasing inter-chromosomal distance. Right, the detachment rate increases with distance, ensuring that distant chromosome pairs form fewer and shorter-lived connections. **(C)** Temporal evolution of inter-chromosomal connectivity for $k_{\text{on}} = 20$ and $k_{\text{off}} = 40$. The fraction of connected chromosome pairs fluctuates over time, maintaining transient connectivity among approximately 20–40% of chromosome pairs at any given moment. **(D)** Experimentally measured displacement correlation as a function of time lag for first-order neighbors in control (Ctl), nocodazole-treated (Noc), and trichostatin A–treated (TSA) cells. Control and Noc-treated cells show similar temporal profiles, with correlations increasing and reaching a comparable plateau. In contrast, TSA-treated cells exhibit a lower plateau and a slower rise in correlation with increasing time lag. Data are shown as mean ± SD. **(E)** Simulated displacement correlation as a function of time lag for first-order neighbors using inter-chromosomal spring parameters calibrated to reproduce the experimentally measured coordination-versus-distance relationships shown in Figure **??**. Simulations capture the reduced correlation amplitude and delayed buildup observed in the TSA condition, but do not reproduce the near-complete overlap between control and Noc curves observed experimentally. Data are shown as mean ± SD.
